# Structural basis of nanobody-recognition of grapevine fanleaf virus and of virus resistance loss

**DOI:** 10.1101/728907

**Authors:** Igor Orlov, Caroline Hemmer, Léa Ackerer, Bernard Lorber, Ahmed Ghannam, Vianney Poignavent, Kamal Hleibieh, Claude Sauter, Corinne Schmitt-Keichinger, Lorène Belval, Jean-Michel Hily, Aurélie Marmonier, Véronique Komar, Sophie Gersch, Pascale Schellenberger, Patrick Bron, Emmanuelle Vigne, Serge Muyldermans, Olivier Lemaire, Gérard Demangeat, Christophe Ritzenthaler, Bruno P. Klaholz

**Affiliations:** Centre for Integrative Biology (CBI), Department of Integrated Structural Biology, IGBMC, Université de Strasbourg, 1 rue Laurent Fries, 67404 Illkirch, France; Institute of Genetics and of Molecular and Cellular Biology (IGBMC), Université de Strasbourg, 1 rue Laurent Fries, Illkirch, France; Centre National de la Recherche Scientifique (CNRS), UMR 7104, Illkirch, France; Institut National de la Santé et de la Recherche Médicale (Inserm), U964, Illkirch, France; Institut de biologie de moléculaire des plantes, UPR2357 du CNRS, Université de Strasbourg, 12 rue du général Zimmer, F-67084 Strasbourg, France; Université de Strasbourg, INRA, SVQV UMR-A 1131, Colmar 68000, France; Institut Français de la Vigne et du Vin, Domaine de l’Espiguette. 30240 Le Grau du Roi, France; Architecture et Réactivité de l’ARN, UPR 9002, Université de Strasbourg, IBMC, CNRS, 15 Rue R. Descartes, 67084 Strasbourg, France; Centre de Biochimie Structurale CNRS INSERM, 29 rue de Navacelles 34090, Montpellier, France; Structural Biology Research Center, Vrije Universiteit Brussel, Pleinlaan 2, 1050 Brussels, Belgium

## Abstract

Grapevine fanleaf virus (GFLV) is a picorna-like plant virus transmitted by nematodes that affects vineyards worldwide. Nanobody (Nb)-mediated resistance against GFLV has been created recently and shown to be highly effective in plants including grapevine, but the underlying mechanism is unknown. Here we present the high-resolution cryo-EM structure of the GFLV-Nb23 complex which provides the basis for the molecular recognition by the nanobody. The structure reveals a composite binding site bridging over 3 domains of the capsid protein (CP) monomer. The structure provides a precise mapping of the Nb23 epitope on the GFLV capsid in which the antigen loop is accommodated through an induced fit mechanism. Moreover, we uncover and characterize several resistance-breaking GFLV isolates with amino acids mapping within this epitope, including C-terminal extensions of the CP, which would sterically interfere with Nb binding. Escape variants with such extended CP fail to be transmitted by nematodes linking Nb-mediated resistance to vector transmission. Together, these data provide insights into the molecular mechanism of Nb23-mediated recognition of GFLV and of virus resistance loss.

**Significance:** Grapevine fanleaf virus (GFLV) is a picorna-like plant virus that severely impacts vineyards worldwide. While Nanobodies (Nb) confer resistance to GFLV in plants the underlying molecular mechanism of action is unknown. Here we present the high-resolution cryo-EM structure of the GFLV-Nb complex. It uncovers the conformational epitope on the capsid surface which is a composite binding site into which the antigen loop is accommodated through an induced fit mechanism. Furthermore, we describe several resistance-breaking isolates of GFLV with reduced Nb binding capacity. Those that carry a C-terminal extension also fail to be transmitted by nematodes. Together, these data provide structure-function insights into the Nb-GFLV recognition and the molecular mechanism leading to loss of resistance.

## Introduction

Vineyards under production for wine grapes, table grapes or dried grapes account for a total of 7.5 million hectares and more than 75.8 million metric tons worldwide for a global trade value of 29 billion Euros (2016 OIV Statistical Report on World Vitiviniculture). This positions grapevine (*Vitis vinifera*) as a highly valuable crop. However, grapevine is susceptible to a wide range of pathogens, including viruses. With over 70 viral species belonging to 27 genera identified so far, grapevine represents the cultivated crop with the highest number of infecting viruses^1^. While the pathogenicity of all these viruses is not established, a number of them cause severe diseases such as the viruses responsible for fanleaf-, leafroll- and rugose wood-diseases that have been reported in nearly all vine growing areas. The grapevine fanleaf virus (GFLV) is a nematode-transmitted virus^13^ and principal causal agent of grapevine fanleaf degeneration. It belongs to the genus *Nepovirus* of the family *Secoviridae* in the order *Picornavirales* and possesses a bipartite single-stranded positive-sense RNA genome^14^. The structure of the icosahedral capsid is known from our previous crystal structure analysis^15^ and follows a pseudo-*T* = 3 triangulation. It is composed of 60 copies of the capsid protein (CP), which folds into three jelly-roll ß sandwiches^15^.

Since their discovery^3^, single-domain antigen-binding fragments of camelid-derived heavy chain-only antibodies, also known as Nanobodies (Nb)^4^, have proven to be of outstanding interest as therapeutics against human diseases and pathogens^5-7^ including viruses^8-10^. Recent reports also revealed their effectiveness to confer resistance against plant viruses. Thus, transient expression of Nb against broad bean mottle virus (BBMV) attenuated the spreading of the cognate virus in *Vicia faba*^11^. Recently, we showed that the constitutive expression of a single Nb named Nb23 specific to GFLV confers resistance to a wide range of GFLV isolates in both the model plant *Nicotiana benthamiana* and grapevine^12^, but the molecular basis of GFLV recognition and the mechanism of resistance induced by Nb23 are unknown. Moreover, while one of the homozygous *N. benthamiana* lines tested was fully resistant to GFLV, another line showed infection at low frequency (3.2%), suggesting the existence of resistance-breaking (RB) variants^12^ whose mechanism of action remains unclear.

To address the molecular basis of Nb23-GFLV recognition, we determined the cryo electron microscopy (cryo-EM) structure of the GFLV-Nb23 complex at high resolution. The structure reveals that Nb23 bridges over 3 domains of the CP and it provides unprecedented insights into the epitope and the residues involved in the interface, including the mechanism of molecular recognition of the antigen-binding loops. We find a perfect correlation between mutations detected in RB variants and the Nb23 epitope observed in the structure, which explains the resistance loss in escape variants (EV). In agreement with the fact that the conformational surface epitope recognized by the Nb23 partially covers a cavity involved in vector transmission^15,16^, we show that EV preexist in natural viral populations at low frequency. We also uncover that the most frequently found EV with extended CP are deficient in transmission by nematodes.

## Results

To gain molecular insights into the mechanism of GFLV recognition by Nb23, precisely map the epitope and decipher the interactions between Nb23 and the CP, we decided to analyze the structure of the GFLV viral particle decorated with Nb23. The structure of the GFLV-Nb23 complex was determined by high-resolution single particle cryo-EM using angular reconstitution^18,19^ and refined to an average resolution of 2.8 Å (Fig. 1; Figs. S1&S2). The cryo-EM map was used to build an atomic model with proper model geometry and statistics using the Phenix software^20^ which involved manual model building and refinement notably of the antigen-binding loop region (see methods). The 3D reconstruction displays the 3 core parts of the complex: the virus capsid, the Nb at the outer surface and the genome inside (Fig. 1). The structure reveals that Nb23 binds at the surface of the GFLV capsid in the vicinity of the 5-fold axis (Fig. 1D-F), at the level of the individual CP monomers that form the pentamers of the pseudo-*T = 3* icosahedral capsid. The outer isocontour surface of the GFLV-Nb23 reconstruction (Fig. 1 and Fig. S1) shows that the Nb23 molecules are positioned far enough from each other to allow 60 of them to attach and reach full 1:1 stoichiometric binding without bridging neighboring CPs. Consistent with this finding, we observe the same stoichiometry by native gel analysis upon titration of GFLV with increasing amounts of Nb23 (Fig. S3).

**Fig. 1.**
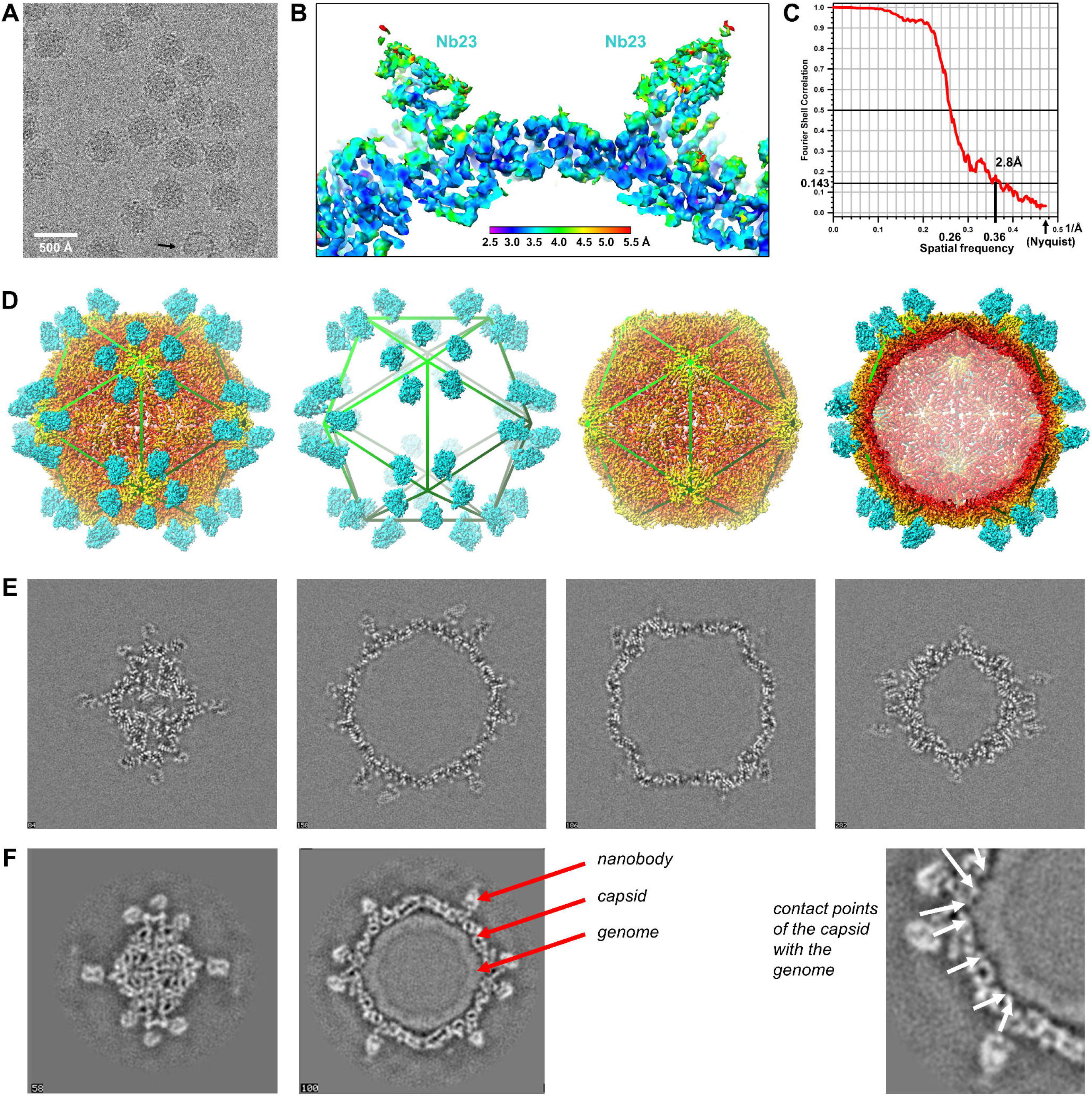
Structural analysis of the GFLV-Nb23 complex by cryo-EM. **A** Typical cryo-EM micrographs of the GFLV-Nb23 complex; most particles are intact and comprise the genome inside (an empty particle is indicated at the bottom with an arrow); scale bar 500 Å). **B** Local resolution estimation, section through Nb23 and the capsid. **C** Plot showing Fourier shell correlation (FSC) versus spatial frequency of the icosahedral averaged reconstruction. Average resolution of the reconstructions is given where the FSC curve crosses below a correlation value of 0.143. **D** Overall structure of the GFLV-Nb23 complex with Nb23 shown in cyan and the capsid in radial coloring (see also Figs. S1&2), including a segmentation of the 3D reconstruction for the Nb23 and the GFLV capsid parts, and a cross section through the icosahedral reconstruction (left to right); the icosahedron is shown as an overlaid polygon. **E** Sections through the refined 3D reconstruction (from top to bottom of the 3D map). **F** Sections through the 3D reconstruction while under refinement; at lower resolution the overall parts are clearly visible: Nb, capsid (secondary structures of the CP are visible) and genome inside the capsid (averaged out due to the icosahedral symmetry that is imposed during 3D reconstruction while the genome might have a different structural organization).

Local resolution estimation shows that while Nb23 is moderately ordered at the periphery (3.5-5 Å resolution towards the C-terminus), it is very well defined in the region of interest enabling a precise analysis of the interactions within the interface between Nb23 and the capsid (Fig. 2). The part of Nb23 that is oriented towards the capsid exhibits clear side-chains densities (Fig. S2) and includes the important antigen-binding loops (Fig. 2D&E). The viral capsid is well defined (including the very N- and C-terminal ends) and comprises the A, B and C domains of each CP (Fig. 2A&B). The interaction region between Nb23 and the capsid involves residues of the complementarity determining regions (CDR) 2 and 3 of Nb23, as well as the two neighboring CP domains A and B (2 out of the 3 jellyroll ß-sandwiches of the CP). These structural elements jointly form a composite binding site with two core interacting regions denoted 1 and 2 (Fig. 2A,D,E), adding up to a total interaction surface of ∼1100 Å^2^. This large binding area is consistent with the formation of a stable Nb23/GFLV complex and with the high affinity of Nb23 for the virus as determined by microscale thermophoresis (Kd = 9.13 ± 2.91 nM, n = 3, Fig. S3).

**Fig. 2.**
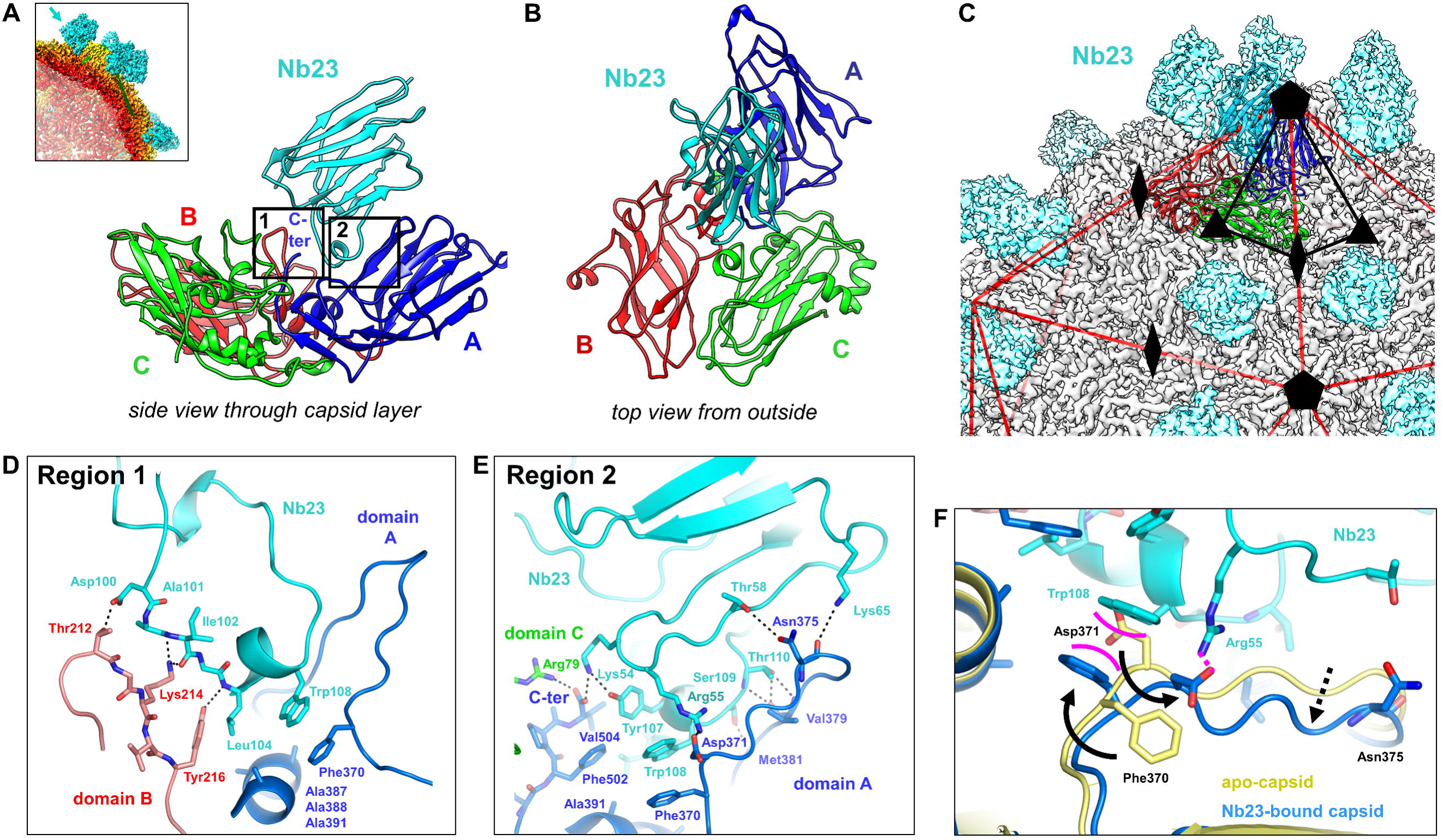
Structural analysis of the Nb23 epitope on the GFLV capsid. **A, B** Side and top view of the CP and Nb23 (cyan), which sits onto a composite binding site comprising the 3 domains of the capsid protein (CP) monomer shown in blue (A), red (B) and green (C); the C-terminus of the CP is indicated. **C** Positions of CP and Nb23 (cyan) with respect to symmetry operators on the icosahedral reconstruction. **D & E** Close-up views of the boxed regions 1 and 2 in panel **A**, showing the detailed molecular interactions (hydrogen bonds are indicated by dotted lines; color code as in panel **B**). **F** Comparison of apo-capsid (in yellow) and Nb23-bound capsid (in blue) showing a conformational adaptation of the capsid residues Phe370 to Asn375, reminiscent of an induced fit mechanism.

The structure allows a precise mapping of the Nb23 epitope on the GFLV capsid and identifies the key residues of Nb23 CDR2 and CDR3 and of GFLV involved in the specific molecular recognition events occurring upon complex formation. Within the two core interacting regions (Fig. 2D,E), a high level of specific interactions is provided through a series of non-covalent bonds (see also Fig. S4). Region 1 (Fig. 2D) comprises loop region 212-216 (ßC’’) of domain B of the CP in which Thr212 interacts with Asp100_Nb23_, Lys214 forms hydrogen bonds with the carbonyl backbone of Ala101_Nb23_ and Ile102_Nb23_, and Tyr216 interacts with the backbone and forms hydrophobic contacts with the side chain of Leu104_Nb23_. Region 2 (Fig. 2E) comprises the two strands of the capsid ß-sheet region 370-391 (domain A of the CP) in which Asp371 forms a salt bridge with Arg55_Nb23_. An additional anchor point of the antibody is provided through Thr58_Nb23_ and Lys65_Nb23_, which interact with Asn375 located in a CP loop (Fig. 2E). In the second ß-strand of the 370-391 region, the backbone of Val379 interacts with Thr110_Nb23_, and the Ser380 backbone and Met381 form contacts with Ser109_Nb23_ and Trp108_Nb23_, respectively. CDR3 forms a long antigen-binding loop with a short α-helical turn carrying Leu104_Nb23_, which is accommodated within a hydrophobic pocket formed by Tyr216, Phe370, Phe502, Val504, and an alanine cluster formed by residues 387, 388 and 391 (Fig. 2D). Trp108_Nb23_ appears to be a key residue in that it inserts into the binding site within a cavity at the junction of the A and B domains of the CP to form π-stacking interactions with Phe370 (Fig. 2E). Importantly, the Phe502 and Val504 residues at the very C-terminal end of the CP are in direct contact with Nb23 and form hydrophobic contacts with Leu104_Nb23_, Tyr107_Nb23_, and Trp108_Nb23_. Finally, Lys54_Nb23_ interact with the free carboxylate moiety of Val504.

Interestingly, comparison of the CP structures in presence and absence of Nb23 (Fig. 2F) reveals that Nb23 induces a conformational change of the CP 370-375 loop region to accommodate the Nb23 side-chains. This significant transition indicates that the molecular recognition of the CP occurs through an induced fit mechanism in which the Phe370 and Asp371 side-chains flip over in opposite directions (Fig. 2F). Without this conformational adaptation, Asp371 would clash with Trp108_Nb23_, which instead needs Phe370 to switch position to form stacking interactions with Trp108, while Asp371 rotates to form a stabilizing salt bridge interaction with Arg55_Nb23_. This selection mechanism may contribute to Nb23 specificity towards the GFLV antigen^12^.

The structure shows that the interface between Nb23 and CP is highly complementary, and also includes the C-terminal CP residue Val504 that is recognized by Lys54_Nb23_. One can therefore hypothesize that disrupting the molecular interface of the epitope region would lead to resistance loss. Interestingly, nanobody-mediated resistance to GFLV in *N. benthamiana* lines that constitutively express Nb23:EGFP displays various degrees of susceptibility to infection indicating that GFLV could overcome Nb-mediated resistance^12^. To further explore this partial resistance breakdown, we forced our two resistant lines towards the emergence of infection events by applying high inoculum pressure (3 μg versus 300 ng of virus). Under such stringent conditions, resistance was indeed overcome by 21 days post inoculation (dpi) in 30 % and 40 % of plants from lines 23EG38-4 and 23EG16-9, respectively (Fig. 3A,B). To address whether the infection events observed were due to the emergence of *bona fide* GFLV RB variants and considering Nb23 is specific to GFLV^12^, we characterized the virus progeny by sequencing the CP from Nb23-expressing plants that showed systemic infection. Remarkably, the analysis reveals that all suspected RB events correspond to viruses with mutations in the CP coding sequence (Fig. 3C). The most frequent mutation (5 independent events) is a nucleotide insertion near the 3’ end of the CP gene that leads to a frameshift responsible for non-conservative mutations of CP residues 503 and 504 and the addition of 12 extra amino acids at the C-terminus of the CP (Fig. 3C). The second most represented mutation (three occurrences) was the suppression of the stop codon leading to a CP with three (CP+3) or five (CP+5) extra C-terminal amino acids, the latter being the consequence of a second mutation. The remaining two mutations are missense mutations mapping to residues 216 (Tyr216His) and 502 (Phe502Leu) of the CP, both of which are directly involved in Nb23-virus recognition (Fig. 2D,E). Remarkably, most RB variants (8 out of 10) display a mutated CP with variable C-terminal extensions of three (CP+3), five or twelve residues (Fig. 3C); such an extension would sterically clash with Nb23 and prevent recognition of the free carboxylate group Val504 as evident from the structure (Fig. 2E). Accordingly, GFLV-CP+3 and GFLV-Tyr_216_His mutants are recognized only very poorly by Nb23 in DAS-ELISA (while they still are with conventional anti-GFLV antibodies; Fig. 3D). To confirm that C-terminal extensions of the CP are RB mutations and to exclude the contribution of non-CP residues in overcoming resistance, the CP+3 mutation was introduced into the GFLV-GHu infectious clone^17^. The resulting mutant fully infects both control and Nb23-expressing plants (100% infection), contrarily to wild-type GFLV-GHu that infects only control plants (Fig. 3E), confirming that the CP+3 mutation is sufficient to overcome Nb-mediated resistance.

**Fig. 3.**
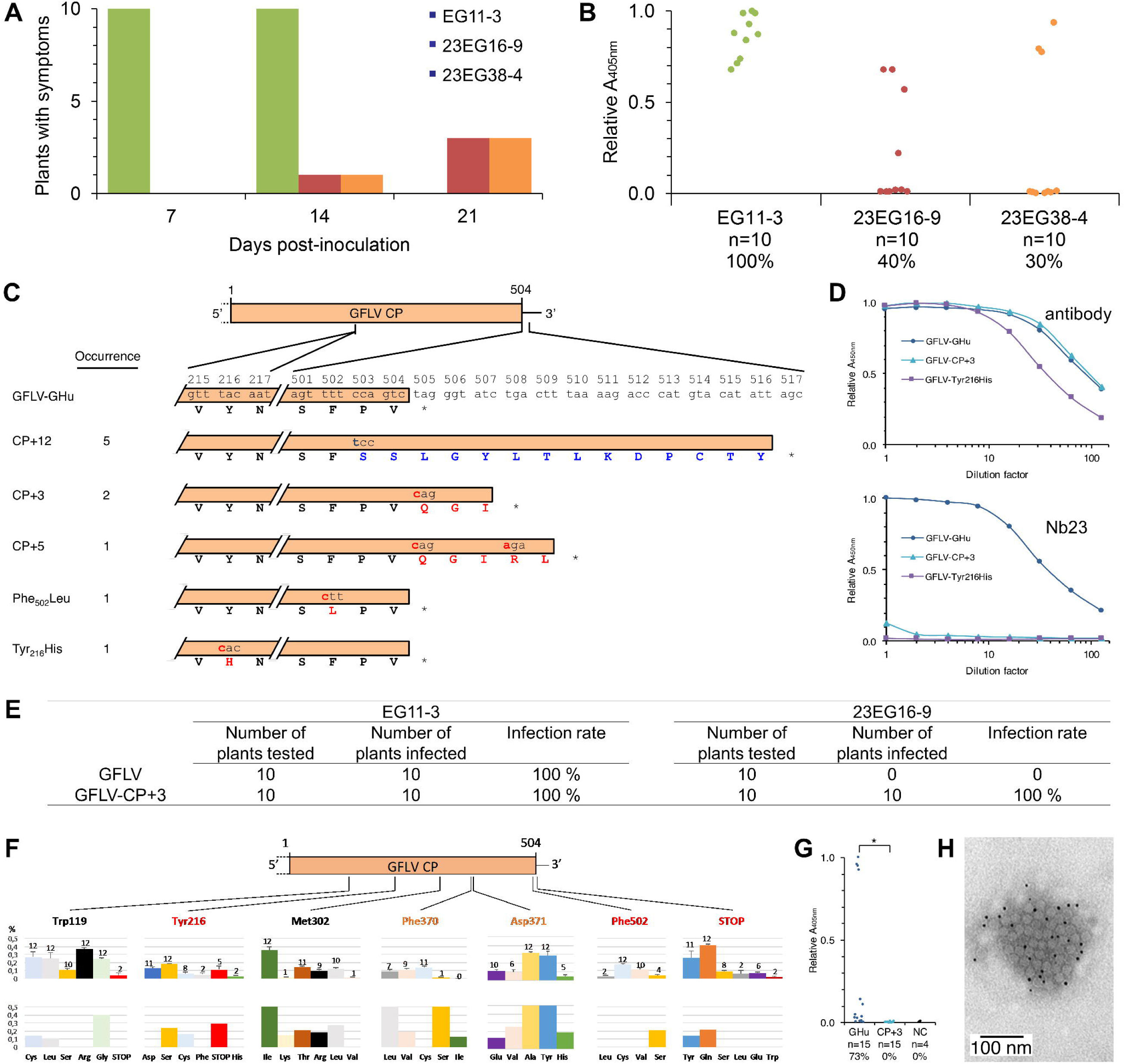
Molecular characterization of EV. **A** Symptom analysis of plants from lines EG11-3 (green), 23EG16-9 (red) and 23EG38-4 (orange) at 7, 14 and 21 dpi upon inoculation with 3 µg of GFLV-GHu. **B** GFLV detection by DAS-ELISA at 21 dpi. Each dot corresponds to a single plant sample and represents the mean relative absorbance at 405 nm of experimental duplicates. Percentage of infections are indicated below each column (%). **C** Mapping of mutations found in EV. Mutations were found in the open reading frame encoding the CP of GFLV (red for substitutions and blue for insertions). In total, sequences from 10 independent EV were obtained and occurrence of mutations is provided for each type of mutant. CP sequence of GFLV-GHu from residues 215 to 217 and 501 to 517 is provided. CP coding sequences are boxed and corresponding protein sequences are provided below. **D** *N. benthamiana* infected with GFLV-GHu, GFLV-CP+3 or GFLV-Tyr_216_His were tested by DAS-ELISA with either conventional antibodies or Nb23 for detection. Conventional antibodies recognize all GFLV isolates contrarily to Nb23 that fails to detect GFLV-CP+3 and GFLV-Tyr_216_His, indicating reduced binding of Nb23 to GFLV EV. **E** Susceptibility of *N. benthamiana* transgenic lines towards GFLV GHu or GFLV-CP+3. Plants were tested by DAS-ELISA for GFLV at 21 dpi. **F** HTS data mining on 13 samples infected with GFLV (12 from grapevines, upper histograms and 1 from nematodes, lower histograms). We focused on seven positions within the CP with residues in red involved in Nb23-CP interactions and found in EV, in orange those involved in Nb23-CP interactions based on the structural analysis (see Fig. 2), and in black those exposed on the outer surface of the capsid but not interacting with Nb23 nor involved in the viral transmission by nematodes. The underlined red residues were found mutated in EV. Each colored bar corresponds to the percentage of a missense mutation occurrence with +/-SE (with the number of event per mutation/site being indicated above). **G** Transmission efficiency of GFLV-GHu and GFLV-CP+3 by nematodes. Dots correspond to DAS-ELISA results from individual bait plants and represent the mean relative absorbance at 405 nm of experimental replicates normalized against maximum assay values. The asterisk indicates statistically significant differences (two tailed paired *t*-test, *P* = 0.022). Number of plants tested (n) and percentage of infections (%) are provided below each column. NC, healthy plants. **H** Negative staining EM image of immunogold-labeled viral particles isolated from GFLV-CP+3 infected plants.

We next investigated the possible pre-existence and frequency of such mutations in natural viral populations from grapevines and nematodes by high throughput screening (HTS) data mining (Fig. 3F, and Table S1), with a focus on the codons encoding Tyr216, Phe502 and the amber stop codon identified to be at the origin of EV (Fig. 3F). We also looked at the codon encoding Phe370 and Asp371 due to the importance of these residues in the interaction with Nb23 as seen in the structure (Fig. 2, see list in Fig. S4). As a control, we analyzed the codons encoding Met302 and Trp119, two outer-exposed residues of the CP excluded from the Nb23 epitope and likely not involved in α/β structures nor in the virus transmission by nematodes^15,16^. All point mutations discovered via Sanger-sequencing were unequivocally identified in our HTS datasets. For example, the Tyr_216_His was found in 2 out of the twelve grapevine samples tested. On the other hand, STOP_505_Gln was detected in all samples at an average rate close to 0.4% together with other missense mutations (*e.g.* STOP_505_Tyr, STOP_505_Ser or STOP_505_Glu), leading to a CP with extra C-terminal amino acid extensions. Such assortment of missense mutations was observed at all examined positions (Fig. 3F). Similarly, most RB mutations were also detected in the viral RNA isolated from viruliferous vector nematodes (Fig. 3F), underscoring that most RB variants preexist in natural viral population at low frequencies as part of the viral quasispecies as defined in Domingo *et al.* ^21^.

Finally, to address the question whether the EV would still be active in vector transmission, we tested GFLV-Tyr_216_His, Phe_502_Leu and CP+3 mutants for transmission by nematodes. While both single point mutants were still vectored, transmission of GFLV-CP+3 was abolished (Fig. 3G) despite its ability to form viral particles (Fig. 3H). This uncovers an important new function of residues in the vicinity of the C-terminal Val504 in GFLV transmission by *Xiphinema index*. Remarkably, as can be seen in the structure, the Nb23 epitope and the ligand-binding pocket (LBP) partially overlap on the capsid surface (Fig. 4A,B). The LBP is predicted to be involved in the specific transmission of GFLV by its nematode vector^15,16^, in particular the loops surrounding the C′C″ loops in the CP B-domains comprising residues Tyr_216_, Phe_502_ and Val_504_ (Fig. 4A,B). The present structure-function analysis thus clarifies why RB mutations not only hinder Nb23 binding but in the case of the C-terminal extensions also interfere with GFLV transmission.

**Fig. 4.**
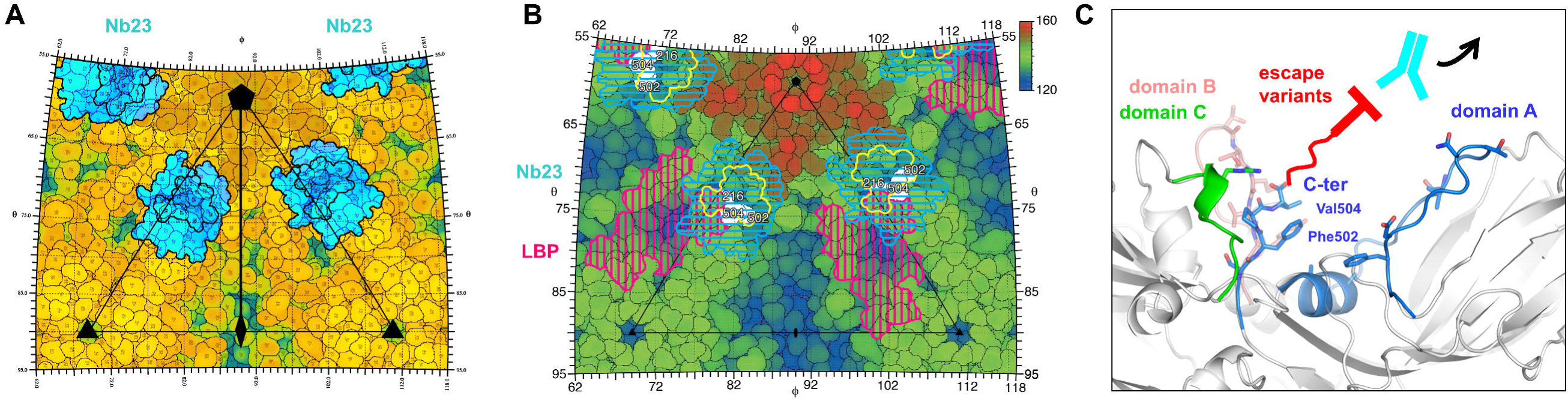
Fingerprint of Nb23 on the GFLV capsid and mechanism of resistance loss. **A** Fingerprint of the Nb23 binding site on the capsid for one icosahedral asymmetric unit (black triangle, see also Fig. S1) in which the polar angles θ and ϕ represent latitude and longitude, respectively. The coloring is according to the radial distance of the surface from the center of the particle. **B** Fingerprint showing the projected surfaces on the GFLV capsid. The Nb23 footprints and projected surfaces are delineated in yellow and cyan, respectively, the ligand binding pockets (LBP) in magenta and mutated residues in GFLV EV shown in white together with amino acid positions. The mosaic background shows the amino acids that form the viral surface. **C** Schematic representation of the molecular mechanism in which the C-terminal extension of GFLV EV would clash with Nb23, thus leading to resistance loss.

Taken together, this study provides new insights into the molecular mechanism of Nb23-mediated recognition of GFLV including the location of the Nb next to the five-fold symmetry axes on the capsid (Fig. 1) and the detailed description of the interface between the GFLV epitope and Nb23 (Fig. 2). The epitope consists of a composite binding site involving domains A, B and C of the CP that interact with the CDR2 and CDR3 of Nb23 (Fig. 2). Remarkably consistent with the epitope seen in the cryo-EM structure, the RB mutations all map to neighboring residues exposed on the outer-surface of the GFLV capsid (Fig. 3 & Fig. S4), which we show preexist at low frequency in virus quasispecies (Fig. 3). Specifically, the structure explains the mechanism of action of the C-terminal extension mutants in overcoming resistance: the extra 3, 5 or 12 residues at the C-terminus of the CP from GFLV EV would interfere with Nb23 binding due to steric clashes and loss of specific hydrogen bonds (in particular between the free carboxylate group of the CP last residue, Val504, and the Lys54_Nb23_ region; Fig. 2E). This would prevent contacts with CP Val504 and occlude the highly specific interface necessary for Nb23-GFLV recognition (Fig. 4C). This manner to interfere with Nb23 binding and prevent GFLV neutralization is likely more potent than single residue missense mutations such as Tyr216His or Phe502Leu, which may explain why most escape mutations consist in C-terminal extensions of the CP. These EV lead to both resistance loss and transmission deficiency due to the overlap between the epitope and the LBP (Fig. 4). Interestingly, this resistance mechanism is different from the one proposed for norovirus-specific Nbs^22^, which is based on an Nb-induced capsid destabilizing activity. However, the specific RB mechanism observed here has its fitness cost because it occurs to the detriment of the transmission of the virus by its vector. In this respect and considering that the RB mutants were isolated from mechanically inoculated *N. benthamiana*, it would be interesting to investigate the emergence of RB isolates also in grapevine upon natural transmission of the virus by nematodes, in particular the possible bias towards the emergence of GFLV isolates with C-terminal extensions of the CP. Their influence on the robustness and durability of the resistance in grapevine under natural conditions remains to be determined especially when aiming at deploying Nb-mediated resistance^12^ as antiviral strategy in vineyards.

## METHODS

### Transgenic *N. benthamiana* lines

Homozygous lines EG11-3 expressing EGFP and 23EG16-9 and 23EG38-4 expressing Nb23 fused to EGFP were described in Hemmer, et al. ^12^.

### Inoculations

Plants were inoculated either mechanically but rub inoculation using carborundum powder or naturally via nematodes. Mechanical inoculations were performed with either purified virus at 300 ng or 3 µg per inoculum or with crude sap from infected plants.

### GFLV DAS-ELISA assessment of Nb23 *in vitro* binding to EV

Frozen leaf samples from infected plants were ground in extraction buffer (35 mM Na_2_HPO_4_, 15 mM KH_2_PO_4_, pH 7.0) in a 1:5 w/v ratio. Virus detection was performed on two-fold serial dilutions of clarified extracts by DAS-ELISA using GFLV antibodies (Bioreba, Catalog # 120275) diluted 1,000-fold in coating buffer (15 mM Na_2_CO_3_, 35 mM NaHCO_3_, pH 9.6) as capture antibody and either GFLV antibodies coupled to alkaline phosphatase (<u>Bioreba,</u> Catalog # 120275) diluted 1,000-fold or Strep II-tagged Nb23 at 1 µg/ml in conjugate buffer (10 mM PBS, 0.1% w/v bovine serum albumin (BSA), 0.05% v/v Tween 20, pH 7.4) as detection antibody. In the latter case, an additional incubation with streptavidin-alkaline phosphatase (Jackson Immunoresearch, Catalog # 016-050-084) at 1:10,000 in conjugate buffer was performed before detection with para-nitrophenyl phosphate (Interchim) at 1 mg/ml in substrate buffer (1 M diethanolamine, pH 9.8). Negative control consisted of non-inoculated healthy plants. Absorbance at 405 nm (A_405nm_) was recorded after one hour of substrate hydrolysis. Results are presented as mean absorbance of experimental duplicates ± standard error normalized against maximum assay value.

### Characterization of RB isolates

Leaf samples from Nb23-expressing *N. benthamiana* that tested positive by ELISA for GFLV were used for immunocapture RT-PCR and gene encoding the CP of GFLV sequenced by Sanger sequencing as described in Martin, et al. ^23^.

### Synthetic GFLV-CP+3 construct

GFLV-CP+3 point mutation was introduced into full-length GFLV-GHu RNA2 cDNA infectious clone (pG2)^17^ as a BglII/SalI fragment following an overlap extension PCR amplification using pG2 as template. GFLV-GHu RNA1, -GHu RNA2 and -CP+3 RNA2 capped transcripts were synthesized from corresponding SalI/XmaI or EcoRI/SalI linearized cDNAs clones by *in vitro* transcription with mMESSAGE mMACHINE T7 kit (Ambion) according to manufacturer’s instructions. For infections, combination of appropriate RNA1 and RNA2 transcripts was used to mechanically inoculate *N. benthamiana* plants at the 4- to 6-leaf stage.

### Transmission assays

Transmission assays by nematodes were performed using a 2-step transmission procedure according to Marmonier, et al. ^24^. During a 6 weeks acquisition access period, ca. 300 aviruliferous *X. index* per plant were left in contact with roots of systemic infected *N. benthamiana* grown in a 3:1:1 v/v ratio of sand, loess and clay pebbles soil. Source plants were then substituted with healthy *N. benthamiana* for a 8 to 10 weeks inoculation access period at the end of which virus transmission was tested on roots of bait plants by DAS-ELISA. GFLV-GHu was used as positive and healthy source plants as negative controls. Values exceeding those of negative controls by a factor of 2.4 were considered positive.

His_6_-tagged and Strep-Tagged II Nb23 were produced as described in Hemmer, et al. ^12^

### Native agarose gel electrophoresis

Native gel electrophoresis of purified GFLV was performed in 1 % w/v agarose gels in 1.0X TA buffer (20 mM TrisBase, 0.06% v/v acetic acid, pH 9). For nucleic-acids detection, 5 µg of virus particles were diluted in loading buffer (10% v/v glycerol, 25 mM HEPES, pH 9) supplemented with ethidium bromide (EtBr) at 0.1 µg/ml. After electrophoretic separation, the EtBr-prestained gel was first processed for nucleic-acid content using the Gel Doc system (Bio-Rad) equipped with a 302 nm excitation source and a 520-640 nm band-pass emission filter before Instant Blue protein staining.

### Microscale thermophoresis (MST)

MST measurements were performed using a Monolith NT115 instrument (NanoTemper Technologies GmbH) using Nb23 and purified GFLV-derived virus-like particles (VLP) in which enhanced GFP is encaged instead of mRFP as described in Belval, et al. ^25^. The concentration of the fluorescent VLP was kept constant at a concentration of 100 nM, while the concentration of Nb23 varied from 0.07 nM to 1.0 μM. Samples were diluted in MST optimized buffer (20 mM Tris-HCL buffer, pH 9.5). Each experiment consisted of 15-point titration series in which the Nb concentrations were generated as a 1:1 dilution of VLP. For the measurements, 10 μL of the different dilutions of samples were filled into hydrophilic treated capillaries (premium coated capillaries, NanoTemper Technologies) and measured after a 10 min equilibration at room temperature. The measurements were performed (n = 3) at 40% LED, and 20 %, 40 %, and 80 % MST power. Laser-On time was 30 s, Laser-Off time 5 s. MST data were analyzed using the program PALMIST (biophysics.swmed.edu/MBR/software.html) and evaluated using the appropriate model (1:1) according to Scheuermann, et al. ^26^.

### Immunosorbent electron microscopy (ISEM)

GFLV-CP+3 infected leaves were ground in phosphate buffer (35 mM Na_2_HPO_4_, 15 mM KH_2_PO_4_), centrifuged at 3,000 x *g* for 5 min and clarified samples incubated for 2 h on carbon-coated Formvar nickel grids (300 mesh, Electron Microscopy Science, Pennsylvania) coated with anti-GFLV polyclonal antibodies (Bioreba AG, Catalog # 120475) at 1:200 dilution. After blocking with 2% w/v BSA, 10% v/v normal goat serum, 0.05% Triton-X100 in 22.5 mM HEPES pH 8 for 30 min, grids were incubated with monoclonal anti-GFLV (Bioreba, Catalog # 120475) at 1:150 dilution for 30 min and immunogold labeling performed using anti-mouse antibodies conjugated to 10 nm colloidal gold particles at 1:50 dilution for 30 min (British Biocell International, Catalog # EM.GAMA10). Experiment was performed at room temperature and washes done with phosphate buffer between all steps. Observations were realized after negative staining with 1% m/v ammonium molybdate using a Philips EM208 transmission electron microscope.

### Sample preparation for cryo-EM

GFLV was purified from infected *N. benthamiana* plants according to Schellenberger, et al. ^15^. GFLV-specific single domain antibody Nanobodies (Nb23) were generated as described in Hemmer, et al. ^12^. Dynamic light scattering analyses were used (i) to find the buffer compositions and pH values at which GFLV and GFLV bound to Nb23 are soluble and (ii) to determine their diameters and monodispersity. Measurements were performed using a Malvern Zetasizer NanoZS instrument and quartz cuvettes. For every 20 µL sample ten successive measurements were done at 20°C with constant amounts of GFLV (0.1 mg/mL). Data were processed with manufacturer’s DTS software (version 6.01). GFLV-F13 remained soluble in 20 mM Tris-HCl pH 8.3 - 8.5 upon addition of Nb23. The latter composition was chosen for the preparation of cryoEM grids in which GFLV at 0.5 mg/ml led to exploitable images.

### Structure determination

Nb23 was mixed with purified GFLV-F13 at a 70:1 or 100:1 molecular ratio so as to exceed the theoretical maximum number of 60 possible binding Nbs per capsid. Samples of the freshly prepared GFLV-Nb23 complex (2.5 µl of a 0.5 mg/mL solution) were applied to 300 mesh holey carbon Quantifoil R1.2/1.3 grids (Quantifoil Micro Tools, Jena, Germany), blotted with filter paper from both sides in the temperature- and humidity-controlled Vitrobot apparatus (FEI, Eindhoven, Netherlands), T = 20 °C, humidity 99%, blot force 4, blot time 0.5 s) and vitrified in liquid ethane pre-cooled by liquid nitrogen. Images were collected on the in-house spherical aberration (Cs) corrected Titan Krios S-FEG instrument (FEI) operating at 300 kV acceleration voltage and at an actual underfocus of Δ□ *z* = −0.6 to −6.0 μm using the second-generation back-thinned direct electron detector CMOS (Falcon II) 4,096 × 4,096 camera and automated data collection with EPU software (FEI, Eindhoven, Netherlands) using seven frames over one second exposure (dose rate of ∼40 e^-^ Å^−2^s^−1^). The calibrated magnification (based on the fit of the GFLV crystal structure into the cryo-EM map) is 127,272× resulting in 0.527 Å pixel size at the specimen level (virtually no difference with the nominal pixel size of 0.533 Å). Before semi-automatic particle picking using IMAGIC^19^, stack alignment was performed using the whole image motion correction method^27^, and correction of the contrast transfer function was done by phase flipping using IMAGIC. After centering by alignment against the total sum reference image, 2D classification in IMAGIC was used to remove bad or empty particles, leaving 6139 out of 9502 particles (selected from 1055 images). For the first steps of image processing, the images were coarsened by 4, and further refinement was achieved with two-times coarsened data. The structure was determined via common lines angle assignment and refined with the anchor set angular reconstitution procedure as implemented in IMAGIC, using final search angles of 0.05°.

The resolution was estimated according to Fourier shell correlation (FSC), indicating an average resolution of 2.8 Å according to the 0.143 and half bit resolution criteria^28,29^. Local resolution values were calculated with ResMap. Map interpretation was done using Chimera^30^ and COOT^31^starting from the available atomic model of the GFLV crystal structure (PDB IDs 4V5T/2Y7T; ^15^) and of a homology model of Nb23 that we derived from the cAb-Lys3 VHH antibody domain using the Swiss-Model Server (swissmodel.expasy.org, starting model: PDB ID 1XFP; ^32^; 68% sequence identity with Nb23). Initial model building was done by a rigid body fitting, followed by extensive manual model building in Coot and real space refinement of the atomic model against the experimental map using Phenix^20^. The final atomic model comprises 295 800 atoms (60 × 4930 atoms excluding hydrogens across the 504 and 137 amino acids for each monomer of the CP and Nb23, respectively). Protein residues of the final atomic model show well-refined geometrical parameters (most favored regions 95.4%, additionally allowed regions 4.6%, and 0.0% of outliers in Ramachandran plots, r.m.s. bond deviations of 0.008Å and angle deviations of 1.3°. Solvent accessible surface was calculated with the program GetArea (probe radius 1.4 A;^33^). Figures were prepared using the software Chimera^30^ and Pymol^34^.

### HTS data mining

To determine whether RB mutations preexisted within natural viral populations or were acquired in Nb23-expressing plants, we analyzed HTS datasets (2×150pb RNAseq performed on an Illumina Hiseq 3000, Illumina, San Diego, CA) from a previous study Hily, et al. ^35^. We specifically focused on 12 repetitions of grafted and non-grafted grapevines naturally infected by a vineyard population of GFLV. In addition, total RNA was extracted from a single sample containing 300 viruliferous nematodes previously fed on GFLV-F13 infected plant according to Demangeat, et al. ^36^ and cDNA library were made as described in Hily, et al. ^35^. Analyses of datasets were performed using CLC Genomics Workbench 8.5.1 software (Qiagen), with mapping parameters set at length fraction of 0.5 and similarity of 0.7. Post-mapping, SNPs at each position were recovered (only SNP above the Illumina error rate of 0.1% per base were recorded) and missense mutations (consisting of 1 SNP per codon) were reported with their percentage rate.

### Statistical analyses

For the GFLV-CP+3 transmission assay, statistical analysis was performed by two-tailed paired *t*-test (*P* = 0,022).

## Acknowledgments

We thank J. Michalon & R. Fritz for IT support, and J.-F. Ménétret for technical support, J. Misbach, L. Ley and the INRA and CNRS greenhouse teams for technical support and plants production and Jean-Yves Sgro (Institute for Molecular Virology, University of Wisconsin, Madison, USA) for helpful advice in stereographic roadmap projections; B.P.K. thanks the late Michael Rossmann for exciting discussions. This work was supported by the CNRS, the Agence Nationale de la Recherche (ANR) (grant ANR-COMBiNiNG ANR-14-CE19-0022), Institut National de la Recherche Agronomique (INRA), Université de Strasbourg, Institut National de la Santé et de la Recherche Médicale (Inserm), the Interreg Rhin Supérieur V project “Vitifutur” http://www.interreg-rhin-sup.eu/projet/vitifutur-reseau-transnational-de-recherche-et-de-formation-en-viticulture/ and the Conseils Interprofessionels des vins d’Alsace, de Champagne, de Bourgogne et de Bordeaux. C.H. was supported by a fellowship from the Région Alsace and the “Plant Health and the Environment” INRA division. L.A. was supported by a CIFRE grant from the Institut Français de la Vigne et du Vin, subsidized by the ANRT (CIFRE convention number 2012/0929). The electron microscope facility at the CBI was supported by the Région Alsace, the Fondation pour la Recherche Médicale (FRM), the IBiSA platform program, Inserm, CNRS, the Association pour la Recherche sur le Cancer (ARC), the French Infrastructure for Integrated Structural Biology (FRISBI) ANR-10-INSB-05-01, and by Instruct-ERIC.

## Contributions

GD, LA, BL & LB prepared samples for cryo-EM; I.O. performed cryo-EM data acquisition, image processing, structure refinement & atomic model building; I.O. & B.P.K. analyzed the structure; CH, VK, AM, EV characterized GFLV escape variants; CS-K cloned the synthetic GFLV-CP+3 construct & performed transcription and inoculation of plants; CH & AM performed the *in vitro* Nb23 affinity assay towards escape variants; LB prepared samples for immunogold labeling by CH; SG, AM, CH & GD analyzed transmission by nematodes; LA, CH & VP produced nanobodies; CH produced transgenic plants and made CP+3 transmission assays and statistical analysis; A.G. & V.P. performed native gel electrophoresis and MST analysis; J-M.H. & OL did HTS, data mining and analysis; B.L. & L.A. did DLS analysis; all authors analyzed the data. B.P.K. and C.R. supervised the project and wrote the manuscript with input from all authors.

## Competing financial interests

The authors declare no competing financial interests.

## Corresponding authors

Correspondence to: Christophe Ritzenthaler or Bruno P. Klaholz

## Accession codes

Cryo-EM map and coordinates of the atomic model can be found in the Protein Data Bank (PDB) and the Electron Microscopy Data Bank (EMD) under the following accession numbers, respectively: PDB (5foj), EMD-3246.

## References

1. Martelli, G. An Overview on Grapevine Viruses, Viroids, and the Diseases They Cause. in Grapevine Viruses: Molecular Biology, Diagnostics and Management 31–46 (Springer, 2017).

2. Oliver, J.E. & Fuchs, M. Tolerance and Resistance to Viruses and Their Vectors in Vitis sp.: A Virologist’s Perspective of the Literature. Am. J. Enol. Vitic. 62, 438–451 (2011).

3. Hamers-Casterman, C. et al. Naturally-occurring antibodies devoid of light-chains. Nature 363, 446–448 (1993).

4. Muyldermans, S. Nanobodies: natural single-domain antibodies. Annu Rev Biochem 82, 775–97 (2013).

5. Kijanka, M., Dorresteijn, B., Oliveira, S. & van Bergen en Henegouwen, P.M. Nanobody-based cancer therapy of solid tumors. Nanomedicine 10, 161–174 (2015).

6. Hassanzadeh-Ghassabeh, G., Devoogdt, N., De Pauw, P., Vincke, C. & Muyldermans, S. Nanobodies and their potential applications. Nanomedicine 8, 1013–1026 (2013).

7. Lo, A.W. et al. The molecular mechanism of Shiga toxin Stx2e neutralization by a single-domain antibody targeting the cell receptor-binding domain. J Biol Chem 289, 25374–81 (2014).

8. Lülf, S. et al. Structural basis for the inhibition of HIV-1 Nef by a high-affinity binding single-domain antibody. Retrovirology 11, 1–13 (2014).

9. Vanlandschoot, P. et al. Nanobodies ® : New ammunition to battle viruses. Antiviral Research 92, 389–407 (2011).

10. Desmyter, A., Spinelli, S., Roussel, A. & Cambillau, C. Camelid nanobodies: killing two birds with one stone. Current Opinion in Structural Biology 32, 1–8 (2015).

11. Ghannam, A., Kumari, S., Muyldermans, S. & Abbady, A.Q. Camelid nanobodies with high affinity for broad bean mottle virus: a possible promising tool to immunomodulate plant resistance against viruses. Plant Mol Biol 87, 355–69 (2015).

12. Hemmer, C. et al. Nanobody-mediated resistance to Grapevine fanleaf virus in plants. Plant Biotechnology Journal 16, 660–671 (2018).

13. Andret-Link, P. et al. Ectoparasitic nematode vectors of grapevine viruses. in Grapevine Viruses: Molecular Biology, Diagnostics and Management 505–529 (Springer, 2017).

14. Schmitt-Keichinger, C., Hemmer, C., Berthold, F. & Ritzenthaler, C. Molecular, Cellular, and Structural Biology of Grapevine fanleaf virus. in Grapevine Viruses: Molecular Biology, Diagnostics and Management (eds. Meng, B., Martelli, G.P., Golino, D.A. & Fuchs, M.) 83–107 (Springer International Publishing, Cham, 2017).

15. Schellenberger, P. et al. Structural insights into viral determinants of nematode mediated Grapevine fanleaf virus transmission. PLoS Pathog 7, e1002034 (2011).

16. Schellenberger, P. et al. A stretch of 11 amino acids in theßB-ßC loop of the coat protein of grapevine fanleaf virus is essential for transmission by the nematode *Xiphinema index*. J. Virol 84, 7924–33 (2010).

17. Vigne, E. et al. A strain-specific segment of the RNA-dependent RNA polymerase of grapevine fanleaf virus determines symptoms in Nicotiana species. J Gen Virol 94, 2803–13 (2013).

18. Van Heel, M. Similarity measures between images. Ultramicroscopy 21, 95–100 (1987).

19. van Heel, M., Harauz, G., Orlova, E.V., Schmidt, R. & Schatz, M. A new generation of the IMAGIC image processing system. Journal of structural biology 116, 17–24 (1996).

20. Afonine, P.V. et al. Towards automated crystallographic structure refinement with phenix. refine. Acta Crystallographica Section D: Biological Crystallography 68, 352–367 (2012).

21. Domingo, E., Sheldon, J. & Perales, C. Viral quasispecies evolution. Microbiology and Molecular Biology Reviews 76, 159–216 (2012).

22. Koromyslova, A.D. & Hansman, G.S. Nanobodies targeting norovirus capsid reveal functional epitopes and potential mechanisms of neutralization. PLoS Pathog 13, e1006636 (2017).

23. Martin, I.R. et al. The 50 distal amino acids of the 2A(HP) homing protein of Grapevine fanleaf virus elicit a hypersensitive reaction on Nicotiana occidentalis. Mol Plant Pathol 19, 731–743 (2018).

24. Marmonier, A. et al. The coat protein determines the specificity of virus transmission by *Xiphinema diversicaudatum*. J. Plant Pathol. 92, 275–279 (2010).

25. Belval, L. et al. Display of whole proteins on inner and outer surfaces of grapevine fanleaf virus-like particles. Plant Biotechnol J 14, 2288–2299 (2016).

26. Scheuermann, T.H., Padrick, S.B., Gardner, K.H. & Brautigam, C.A. On the acquisition and analysis of microscale thermophoresis data. Analytical biochemistry 496, 79–93 (2016).

27. Li, X. et al. Electron counting and beam-induced motion correction enable near-atomic-resolution singleparticle cryo-EM. Nature methods 10, 584 (2013).

28. Rosenthal, P.B. & Henderson, R. Optimal determination of particle orientation, absolute hand, and contrast loss in single-particle electron cryomicroscopy. Journal of molecular biology 333, 721–745 (2003).

29. Van Heel, M. & Schatz, M. Fourier shell correlation threshold criteria. Journal of structural biology 151, 250–262 (2005).

30. Pettersen, E.F. et al. UCSF Chimera - A visualization system for exploratory research and analysis. J. Comput. Chem. 25, 1605–1612 (2004).

31. Emsley, P., Lohkamp, B., Scott, W.G. & Cowtan, K. Features and development of Coot. Acta Crystallographica Section D: Biological Crystallography 66, 486–501 (2010).

32. De Genst, E. et al. Chemical basis for the affinity maturation of a camel single domain antibody. Journal of Biological Chemistry 279, 53593–53601 (2004).

33. Fraczkiewicz, R. & Braun, W. Exact and efficient analytical calculation of the accessible surface areas and their gradients for macromolecules. Journal of computational chemistry 19, 319–333 (1998).

34. DeLano, W.L. Pymol: An open-source molecular graphics tool. CCP4 Newsletter On Protein Crystallography 40, 82–92 (2002).

35. Hily, J.-M. et al. High-Throughput Sequencing and the Viromic Study of Grapevine Leaves: From the Detection of Grapevine-Infecting Viruses to the Description of a New Environmental Tymovirales Member. Frontiers in Microbiology 9(2018).

36. Demangeat, G., Komar, V., Cornuet, P., Esmenjaud, D. & Fuchs, M. Sensitive and reliable detection of grapevine fanleaf virus in a single Xiphinema index nematode vector. J Virol Methods 122, 79–86 (2004).

