## Supplementary Figures for "Structural basis of nanobody-recognition of grapevine fanleaf virus and of virus resistance loss"

### Supplementary Data

A

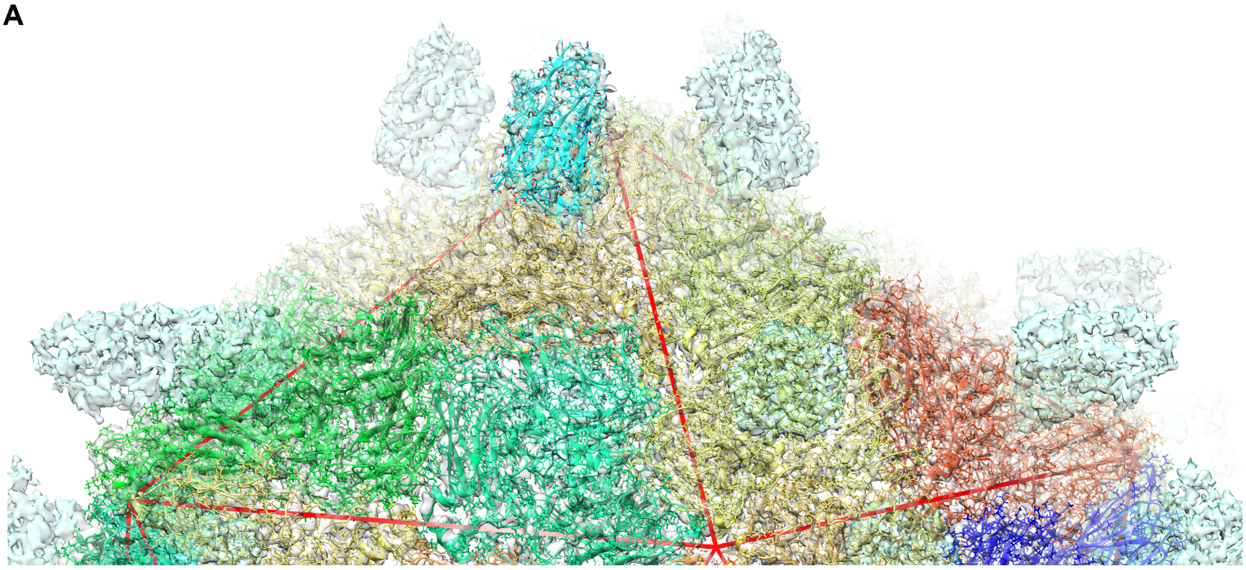

B

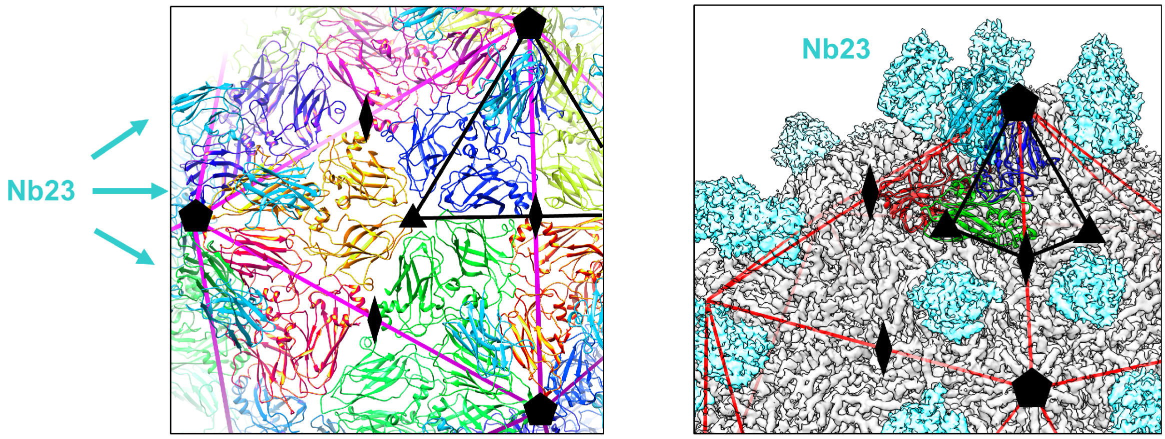


*coloured by CP monomers; top view*

C

| Data collection | |
|---|---|
| **Microscope** | Titan Krios |
| **Acceleration voltage** | 300 kV |
| **Magnification** | 120,000 times |
| **Nominal pixel size** | 0.533 Å / pixel |
| **Actual pixel size** | 0.527 Å / pixel |
| **Detector camera** | Direct electron detector (Falcon II camera) |
| **Detector mode** | Movie mode |
| **Number of movie frames** | 7 |
| **Exposure time** | 1 sec |
| **Electron dose** | ~40 electrons / Å² / sec |
| **Number of images** | 2271 |
| **Number of images used** | 1055 |
| **Closest to focus image** | ~ -0.6 microns |
| **Furthest from focus image** | ~ -6.0 microns |
| **Number of particles selected** | 9502 |
| **Number of particles used** | 6139 |

D

| **Coarsening (times or pixels)** | **4** | **2** | **non-coarsened** |
|---|---|---|---|
| **Nominal pixel size (Å)** | 2.108 (2.132) | 1.054 (1.066) | 0.527 (0.533) |
| **Nyquist-Shannon limit (Å)** | 8.416 (8.528) | 4.208 (4.264) | 2.104 (2.132) |

E

| **Atomic model composition for an asymmetric unit** | |
|---|---|
| Non-hydrogen atoms | 4929 |
| Amino acids | 634 |
| **Atomic model statistics** | |
| Ramachandran favoured | 95.4 % |
| Ramachandran allowed | 4.6 % |
| Ramachandran outliers | 0.0 % |
| All-atom clash score | 5.42 |
| Rotamer outliers | 1.7 % |
| C-beta deviations | 0 |

**Structural environment and symmetries in the GFLV-Nb23 complex & structure data statistics**

**A** View on the entire complex. **B** Positions of CP and Nb23 (cyan) with respect to symmetry operators on the icosahedral reconstruction (the CP monomers are colored individually in the atomic model on the left, and in grey in the cryo-EM map on the right). **C** Cryo-EM data collection and atomic model refinement parameters.

**Supplementary Fig. S1**

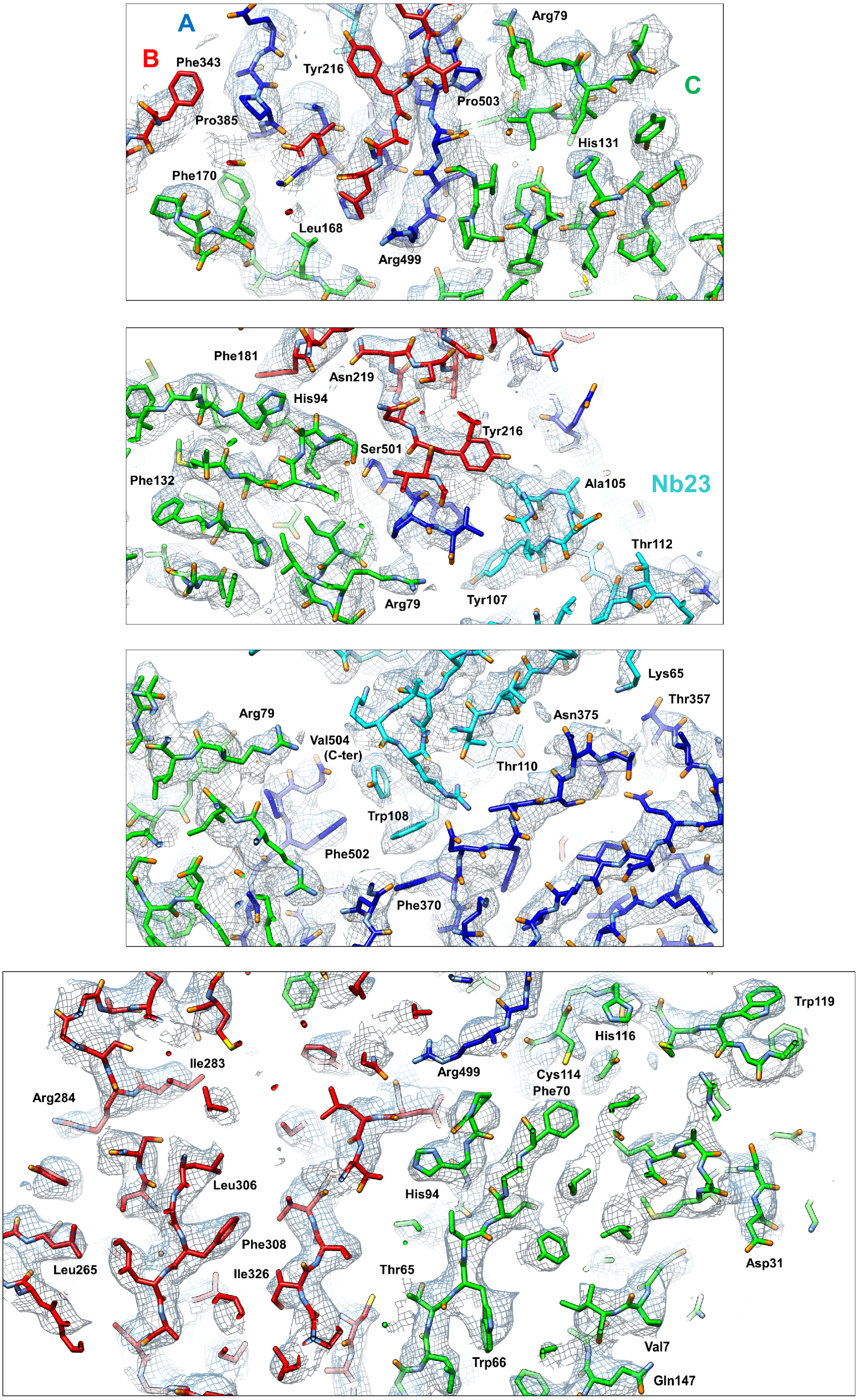


**Details of the cryo-EM map and atomic model of the GFLV-Nb23 complex**

Top to bottom, detailed views of representative regions of the cryo-EM map and the refined atomic model.

**Supplementary Fig. S2**

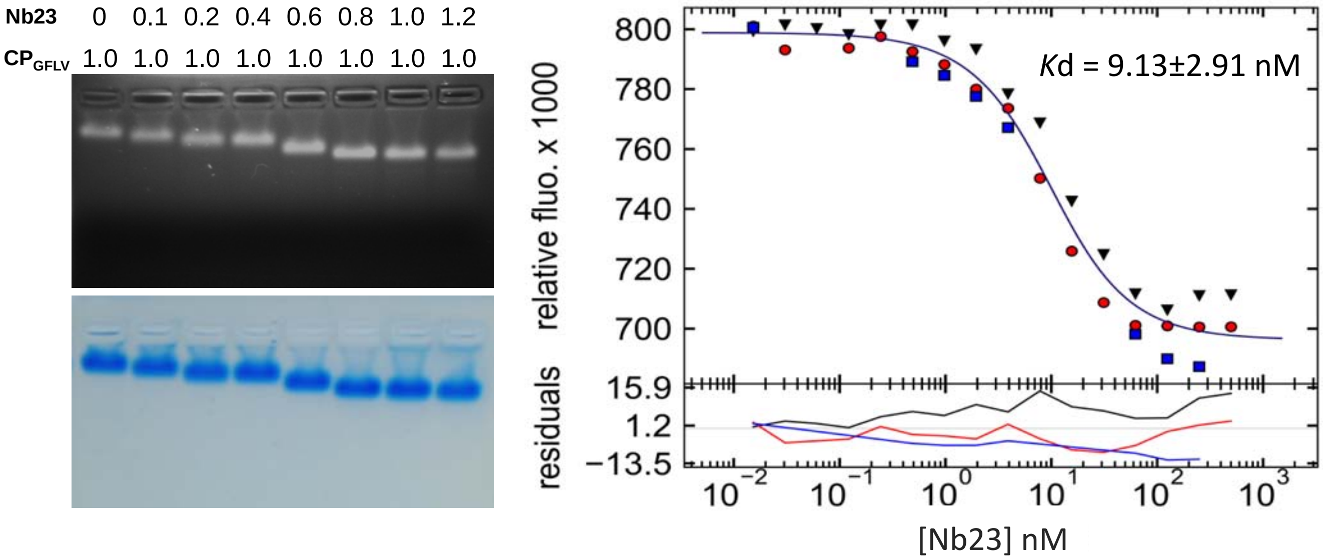


**Stoichiometry determined by native gel electrophoresis & binding of Nb23 to GFP-labeled GFLV-derived VLP analyzed by MicroScale Thermophoresis (MST)**

*Left* Native agarose gel electrophoresis of GFLV decorated with Nb23. GFLV (5 µg/lane) was incubated with increasing amounts of Nb23 (from 0 to 1.7 µg/lane) before electrophoresis. Molar ratios between Nb23 and GFLV CP are given above each lane. Upper panel: fluorescence detection of GFLV after ethidium bromide staining and UV illumination. Lower panel: same gel after instant-blue staining. *Right* The binding curve is derived from the specific change in the thermophoretic mobility upon Nb23 titration (0.07 nM - 10 µM) to a constant concentration (100 nM) of fluorescently labeled VLP. Analysis was performed in triplicate indicated by three colors on the upper plot (11 reading points per color). Fitted binding curve is overlaid onto data points of triplicate. Relative Fluorescence represents ratio (expressed in per-mille units) of the fluorescence readings in the time traces.

**Supplementary Fig. S3**

<table>
<tr><th>Nb23 residues</th><th>CP residues</th><th>Contact type</th><th>CDR</th><th>CP domain</th></tr>
<tr><td>Lys54 side-chain</td><td>Val504 terminal carboxylate</td><td>salt bridge</td><td>2</td><td rowspan="4">A</td></tr>
<tr><td>Arg55 side-chain</td><td>Asp371 side-chain</td><td>salt bridge</td><td>2</td></tr>
<tr><td>Thr58 side-chain</td><td>Asn375 side-chain</td><td>hydrogen bond</td><td>2</td></tr>
<tr><td>Lys65 side-chain</td><td>Asn375 backbone</td><td>hydrogen bond</td><td>2</td></tr>
<tr><td>Asp100 side-chain</td><td>Thr212 side-chain</td><td>hydrogen bond</td><td>3</td><td rowspan="3">B</td></tr>
<tr><td>Ala101 backbone</td><td>Lys214 side-chain</td><td>hydrogen bond</td><td>3</td></tr>
<tr><td>Ile102 backbone</td><td>Lys214 side-chain</td><td>hydrogen bond</td><td>3</td></tr>
<tr><td>Leu104 backbone</td><td>Tyr216 side-chain</td><td>hydrogen bond</td><td>3</td><td>A/B</td></tr>
<tr><td>Leu104 side-chain</td><td>Tyr216 side-chain</td><td>hydrophobic contact</td><td>3</td><td>A/B</td></tr>
<tr><td>Leu104 side-chain</td><td>Ala387 side-chain<br>Ala388 side-chain<br>Ala391 side-chain<br>Phe502 side-chain<br>Val504 side-chain</td><td>hydrophobic contact</td><td>3</td><td rowspan="6">A</td></tr>
<tr><td>Tyr107 side-chain</td><td>Phe502 side-chain<br>Val504 side-chain</td><td>hydrophobic contact</td><td>3</td></tr>
<tr><td>Trp108 side-chain</td><td>Phe370 side-chain</td><td>π-stacking</td><td>3</td></tr>
<tr><td>Trp108 side-chain</td><td>Asp371 side-chain<br>Ala391 side-chain<br>Phe502 side-chain<br>Met381 side-chain</td><td>hydrophobic contact</td><td>3</td></tr>
<tr><td>Ser109 side-chain</td><td>Met381 backbone</td><td>hydrogen bond</td><td>3</td></tr>
<tr><td>Thr110 side-chain and backbone</td><td>Val379 backbone</td><td>hydrogen bonds</td><td>3</td></tr>
</table>

**List of amino acids in the GFLV-Nb23 complex at the interface of the capsid and the nanobody**
The involved residues of the Nb23 nanobody and the GFLV capsid protein are indicated, together with the contact type (hydrogen bond, hydrophobic contact etc.), the CDR and CP regions.

**Supplementary Fig. S4**
