## Supplementary Table 1 for "Structural basis of nanobody-recognition of grapevine fanleaf virus and of virus resistance loss"

| Technic | Sample type | Sample name | codon position | Trp (W) 119 | | | | | aa | % occurrence |
|---|---|---|---|---|---|---|---|---|---|---|
| RNAseq 2*150 | natural population | WTR1 | 1 | T | A | C | G | | W | |
| | | | | 56510 | 112 | 125 | 103 | 56850 | C | 0.903 |
| | | | 2 | T | A | C | G | | L | 1.000 |
| | | | | 570 | 75 | 77 | 56288 | 57010 | S | 0.135 |
| | | | 3 | T | A | C | G | | R | 0.417 |
| | | | | 458 | 64 | 65 | 57363 | 57950 | G | 0.181 |
| | | | | | | | | | STOP | 0.242 |
| RNAseq 2*150 | natural population | WTR2 | 1 | T | A | C | G | | W | |
| | | | | 40145 | 47 | 87 | 96 | 40375 | C | 0.108 |
| | | | 2 | T | A | C | G | | L | 0.138 |
| | | | | 56 | 31 | 50 | 40350 | 40487 | S | 0.123 |
| | | | 3 | T | A | C | G | | R | 0.332 |
| | | | | 31 | 26 | 44 | 40731 | 40832 | G | 0.238 |
| | | | | | | | | | STOP | 0.000 |
| RNAseq 2*150 | natural population | WTR3 | 1 | T | A | C | G | | W | |
| | | | | 47989 | 50 | 121 | 110 | 48270 | C | 0.227 |
| | | | 2 | T | A | C | G | | L | 0.167 |
| | | | | 81 | 27 | 66 | 48245 | 48419 | S | 0.136 |
| | | | 3 | T | A | C | G | | R | 0.354 |
| | | | | 57 | 29 | 54 | 48809 | 48949 | G | 0.228 |
| | | | | | | | | | STOP | 0.000 |
| RNAseq 2*150 | natural population | WTR4 | 1 | T | A | C | G | | W | |
| | | | | 29655 | 64 | 63 | 32 | 29814 | C | 0.535 |
| | | | 2 | T | A | C | G | | L | 0.451 |
| | | | | 135 | 36 | 25 | 29725 | 29921 | S | 0.000 |
| | | | 3 | T | A | C | G | | R | 0.426 |
| | | | | 127 | 33 | 35 | 30104 | 30299 | G | 0.107 |
| | | | | | | | | | STOP | 0.229 |
| RNAseq 2*150 | natural population | WTR5 | 1 | T | A | C | G | | W | |
| | | | | 43208 | 46 | 111 | 109 | 43474 | C | 0.102 |
| | | | 2 | T | A | C | G | | L | 0.128 |
| | | | | 56 | 23 | 49 | 43498 | 43626 | S | 0.112 |
| | | | 3 | T | A | C | G | | R | 0.361 |
| | | | | 29 | 22 | 45 | 43967 | 44063 | G | 0.251 |
| | | | | | | | | | STOP | 0.000 |
| RNAseq 2*150 | natural population | WTR6 | 1 | T | A | C | G | | W | |
| | | | | 30670 | 37 | 71 | 81 | 30859 | C | 0.137 |
| | | | 2 | T | A | C | G | | L | 0.187 |
| | | | | 58 | 13 | 33 | 30877 | 30981 | S | 0.107 |
| | | | 3 | T | A | C | G | | R | 0.350 |
| | | | | 21 | 27 | 43 | 31209 | 31300 | G | 0.262 |
| | | | | | | | | | STOP | 0.000 |
| RNAseq 2*150 | natural population | SWT1 | 1 | T | A | C | G | | W | |
| | | | | 37610 | 42 | 95 | 88 | 37835 | C | 0.231 |
| | | | 2 | T | A | C | G | | L | 0.145 |
| | | | | 55 | 30 | 56 | 37769 | 37910 | S | 0.148 |
| | | | 3 | T | A | C | G | | R | 0.362 |
| | | | | 48 | 18 | 41 | 38452 | 38559 | G | 0.233 |
| | | | | | | | | | STOP | 0.000 |
| RNAseq 2*150 | natural population | SWT2 | 1 | T | A | C | G | | W | |
| | | | | 25366 | 45 | 64 | 75 | 25550 | C | 0.227 |
| | | | 2 | T | A | C | G | | L | 0.206 |
| | | | | 53 | 15 | 35 | 25580 | 25683 | S | 0.136 |
| | | | 3 | T | A | C | G | | R | 0.427 |
| | | | | 32 | 13 | 27 | 25966 | 26038 | G | 0.294 |
| | | | | | | | | | STOP | 0.000 |
| RNAseq 2*150 | natural population | SWT3 | 1 | T | A | C | G | | W | |
| | | | | 16441 | 20 | 42 | 46 | 16549 | C | 0.131 |
| | | | 2 | T | A | C | G | | L | 0.108 |
| | | | | 18 | 13 | 17 | 16560 | 16608 | S | 0.102 |
| | | | 3 | T | A | C | G | | R | 0.375 |
| | | | | 14 | 8 | 22 | 16717 | 16761 | G | 0.278 |
| | | | | | | | | | STOP | 0.000 |
| RNAseq 2*150 | natural population | SWT4 | 1 | T | A | C | G | | W | |
| | | | | 7947 | 6 | 15 | 22 | 7990 | C | 0.110 |
| | | | 2 | T | A | C | G | | L | 0.137 |
| | | | | 11 | 7 | 6 | 7992 | 8016 | S | 0.000 |
| | | | 3 | T | A | C | G | | R | 0.188 |
| | | | | 8 | 4 | 9 | 8145 | 8166 | G | 0.275 |
| | | | | | | | | | STOP | 0.000 |
| RNAseq 2*150 | natural population | SWT5 | 1 | T | A | C | G | | W | |
| | | | | 8717 | 18 | 22 | 26 | 8783 | C | 0.359 |
| | | | 2 | T | A | C | G | | L | 0.125 |
| | | | | 11 | 6 | 12 | 8786 | 8815 | S | 0.136 |
| | | | 3 | T | A | C | G | | R | 0.455 |
| | | | | 11 | 5 | 21 | 8874 | 8911 | G | 0.296 |
| | | | | | | | | | STOP | 0.000 |
| RNAseq 2*150 | natural population | SWT6 | 1 | T | A | C | G | | W | |
| | | | | 20473 | 21 | 46 | 50 | 20590 | C | 0.129 |
| | | | 2 | T | A | C | G | | L | 0.189 |
| | | | | 39 | 15 | 23 | 20572 | 20649 | S | 0.111 |
| | | | 3 | T | A | C | G | | R | 0.325 |
| | | | | 11 | 11 | 27 | 20877 | 20926 | G | 0.243 |
| | | | | | | | | | STOP | 0.000 |
| RNAseq 2*150 | natural F13 pop. | mix of 300 | 1 | T | A | C | G | | W | |
| | | nematods | | 2715 | 1 | 2 | 11 | 2729 | C | 0.143 |
| | | | 2 | T | A | C | G | | L | 0.110 |
| | | | | 3 | 1 | 1 | 2732 | 2737 | S | 0.000 |
| | | | 3 | T | A | C | G | | R | 0.000 |
| | | | | 4 | 0 | 0 | 2789 | 2793 | G | 0.403 |
| | | | | | | | | | STOP | 0.000 |

| Sample name | codon position | Tyr (Y) 216 | | | | | aa | % occurrence |
|---|---|---|---|---|---|---|---|---|
| WTR1 | 1 | T | A | C | G | | Y | |
| | | 56596 | 37 | 50 | 66 | 56749 | D | 0.116 |
| | 2 | T | A | C | G | | S | 0.176 |
| | | 164 | 55875 | 99 | 61 | 56199 | C | 0.109 |
| | 3 | T | A | C | G | | F | 0.292 |
| | | 11243 | 189 | 44388 | 78 | 55898 | STOP | 0.478 |
| | | | | | | | H | 0.000 |
| WTR2 | 1 | T | A | C | G | | Y | |
| | | 44586 | 9 | 36 | 69 | 44700 | D | 0.154 |
| | 2 | T | A | C | G | | S | 0.202 |
| | | 24 | 43856 | 89 | 41 | 44010 | C | 0.000 |
| | 3 | T | A | C | G | | F | 0.000 |
| | | 57 | 55 | 43670 | 39 | 43821 | STOP | 0.126 |
| | | | | | | | H | 0.000 |
| WTR3 | 1 | T | A | C | G | | Y | |
| | | 55757 | 9 | 50 | 79 | 55895 | D | 0.141 |
| | 2 | T | A | C | G | | S | 0.225 |
| | | 49 | 55003 | 124 | 57 | 55233 | C | 0.103 |
| | 3 | T | A | C | G | | F | 0.089 |
| | | 19577 | 45 | 35292 | 54 | 54968 | STOP | 0.000 |
| | | | | | | | H | 0.000 |
| WTR4 | 1 | T | A | C | G | | Y | |
| | | 33643 | 13 | 20 | 35 | 33711 | D | 0.104 |
| | 2 | T | A | C | G | | S | 0.162 |
| | | 69 | 33202 | 54 | 44 | 33369 | C | 0.132 |
| | 3 | T | A | C | G | | F | 0.000 |
| | | 3415 | 94 | 29685 | 50 | 33244 | STOP | 0.433 |
| | | | | | | | H | 0.000 |
| WTR5 | 1 | T | A | C | G | | Y | |
| | | 48316 | 7 | 38 | 71 | 48432 | D | 0.147 |
| | 2 | T | A | C | G | | S | 0.203 |
| | | 30 | 47667 | 97 | 64 | 47858 | C | 0.134 |
| | 3 | T | A | C | G | | F | 0.000 |
| | | 24076 | 31 | 23594 | 45 | 47746 | STOP | 0.000 |
| | | | | | | | H | 0.000 |
| WTR6 | 1 | T | A | C | G | | Y | |
| | | 33866 | 6 | 25 | 38 | 33935 | D | 0.112 |
| | 2 | T | A | C | G | | S | 0.162 |
| | | 14 | 33279 | 54 | 28 | 33375 | C | 0.000 |
| | 3 | T | A | C | G | | F | 0.000 |
| | | 45 | 22 | 33082 | 20 | 33169 | STOP | 0.000 |
| | | | | | | | H | 0.000 |
| SWT1 | 1 | T | A | C | G | | Y | |
| | | 36200 | 15 | 25 | 40 | 36280 | D | 0.110 |
| | 2 | T | A | C | G | | S | 0.215 |
| | | 14 | 35752 | 77 | 34 | 35877 | C | 0.000 |
| | 3 | T | A | C | G | | F | 0.000 |
| | | 82 | 51 | 35449 | 26 | 35608 | STOP | 0.216 |
| | | | | | | | H | 0.000 |
| SWT2 | 1 | T | A | C | G | | Y | |
| | | 25408 | 7 | 31 | 34 | 25480 | D | 0.133 |
| | 2 | T | A | C | G | | S | 0.158 |
| | | 20 | 25164 | 40 | 23 | 25247 | C | 0.091 |
| | 3 | T | A | C | G | | F | 0.000 |
| | | 11336 | 20 | 13745 | 25 | 25126 | STOP | 0.000 |
| | | | | | | | H | 0.000 |
| SWT3 | 1 | T | A | C | G | | Y | |
| | | 17517 | 3 | 18 | 16 | 17554 | D | 0.000 |
| | 2 | T | A | C | G | | S | 0.150 |
| | | 9 | 17255 | 26 | 12 | 17302 | C | 0.000 |
| | 3 | T | A | C | G | | F | 0.000 |
| | | 5315 | 17 | 11891 | 15 | 17238 | STOP | 0.000 |
| | | | | | | | H | 0.103 |
| SWT4 | 1 | T | A | C | G | | Y | |
| | | 8326 | 0 | 8 | 18 | 8352 | D | 0.216 |
| | 2 | T | A | C | G | | S | 0.157 |
| | | 3 | 8260 | 13 | 9 | 8285 | C | 0.109 |
| | 3 | T | A | C | G | | F | 0.000 |
| | | 17 | 3 | 8203 | 6 | 8229 | STOP | 0.000 |
| | | | | | | | H | 0.000 |
| SWT5 | 1 | T | A | C | G | | Y | |
| | | 9236 | 4 | 10 | 14 | 9264 | D | 0.151 |
| | 2 | T | A | C | G | | S | 0.164 |
| | | 4 | 9143 | 15 | 8 | 9170 | C | 0.087 |
| | 3 | T | A | C | G | | F | 0.000 |
| | | 2454 | 5 | 6651 | 12 | 9122 | STOP | 0.186 |
| | | | | | | | H | 0.108 |
| SWT6 | 1 | T | A | C | G | | Y | |
| | | 20633 | 1 | 19 | 27 | 20680 | D | 0.131 |
| | 2 | T | A | C | G | | S | 0.196 |
| | | 7 | 20341 | 40 | 27 | 20415 | C | 0.132 |
| | 3 | T | A | C | G | | F | 0.000 |
| | | 793 | 20 | 19462 | 14 | 20289 | STOP | 0.000 |
| | | | | | | | H | 0.000 |
| mix of 300 nematods | 1 | T | A | C | G | | Y | |
| | | 3791 | 1 | 0 | 3 | 3795 | D | 0.295 |
| | 2 | T | A | C | G | | S | 0.239 |
| | | 2 | 3751 | 9 | 6 | 3768 | C | 0.159 |
| | 3 | T | A | C | G | | F | 0.000 |
| | | 3711 | 1 | 8 | 11 | 3731 | STOP | 0.000 |
| | | | | | | | H | 0.000 |

| Sample name | codon position | Met (M) 302 | | | | | aa | % occurrence |
|---|---|---|---|---|---|---|---|---|
| WTR1 | 1 | T | A | C | G | | M | |
| | | 114 | 64505 | 43 | 60 | 64722 | I | 0.715 |
| | 2 | T | A | C | G | | K | 0.000 |
| | | 64004 | 60 | 80 | 69 | 64213 | T | 0.125 |
| | 3 | T | A | C | G | | R | 0.107 |
| | | 245 | 107 | 109 | 63985 | 64446 | L | 0.176 |
| | | | | | | | V | 0.000 |
| WTR2 | 1 | T | A | C | G | | M | |
| | | 6 | 41799 | 39 | 30 | 41874 | I | 0.315 |
| | 2 | T | A | C | G | | K | 0.000 |
| | | 41393 | 28 | 55 | 31 | 41507 | T | 0.133 |
| | 3 | T | A | C | G | | R | 0.000 |
| | | 59 | 31 | 72 | 41456 | 41618 | L | 0.000 |
| | | | | | | | V | 0.000 |
| WTR3 | 1 | T | A | C | G | | M | |
| | | 5 | 38440 | 44 | 19 | 38508 | I | 0.322 |
| | 2 | T | A | C | G | | K | 0.000 |
| | | 38031 | 31 | 60 | 50 | 38172 | T | 0.157 |
| | 3 | T | A | C | G | | R | 0.131 |
| | | 37 | 30 | 86 | 38104 | 38257 | L | 0.114 |
| | | | | | | | V | 0.000 |
| WTR4 | 1 | T | A | C | G | | M | |
| | | 46 | 33199 | 25 | 34 | 33304 | I | 0.551 |
| | 2 | T | A | C | G | | K | 0.000 |
| | | 32900 | 24 | 42 | 39 | 33005 | T | 0.127 |
| | 3 | T | A | C | G | | R | 0.118 |
| | | 95 | 47 | 40 | 32861 | 33043 | L | 0.138 |
| | | | | | | | V | 0.102 |
| WTR5 | 1 | T | A | C | G | | M | |
| | | 8 | 38293 | 49 | 31 | 38381 | I | 0.274 |
| | 2 | T | A | C | G | | K | 0.000 |
| | | 38015 | 28 | 69 | 55 | 38167 | T | 0.181 |
| | 3 | T | A | C | G | | R | 0.144 |
| | | 38 | 36 | 67 | 38133 | 38274 | L | 0.128 |
| | | | | | | | V | 0.000 |
| WTR6 | 1 | T | A | C | G | | M | |
| | | 6 | 34701 | 50 | 29 | 34786 | I | 0.257 |
| | 2 | T | A | C | G | | K | 0.000 |
| | | 34399 | 13 | 55 | 46 | 34513 | T | 0.159 |
| | 3 | T | A | C | G | | R | 0.133 |
| | | 32 | 18 | 57 | 34506 | 34613 | L | 0.144 |
| | | | | | | | V | 0.000 |
| SWT1 | 1 | T | A | C | G | | M | |
| | | 12 | 41929 | 47 | 32 | 42020 | I | 0.266 |
| | 2 | T | A | C | G | | K | 0.000 |
| | | 41514 | 21 | 37 | 43 | 41615 | T | 0.000 |
| | 3 | T | A | C | G | | R | 0.103 |
| | | 44 | 23 | 67 | 41658 | 41792 | L | 0.112 |
| | | | | | | | V | 0.000 |
| SWT2 | 1 | T | A | C | G | | M | |
| | | 3 | 24969 | 32 | 17 | 25021 | I | 0.469 |
| | 2 | T | A | C | G | | K | 0.101 |
| | | 24724 | 25 | 38 | 38 | 24825 | T | 0.153 |
| | 3 | T | A | C | G | | R | 0.153 |
| | | 34 | 27 | 56 | 24813 | 24930 | L | 0.128 |
| | | | | | | | V | 0.000 |
| SWT3 | 1 | T | A | C | G | | M | |
| | | 2 | 16789 | 23 | 10 | 16824 | I | 0.335 |
| | 2 | T | A | C | G | | K | 0.000 |
| | | 16619 | 10 | 28 | 21 | 16678 | T | 0.168 |
| | 3 | T | A | C | G | | R | 0.126 |
| | | 17 | 9 | 39 | 16669 | 16734 | L | 0.137 |
| | | | | | | | V | 0.000 |
| SWT4 | 1 | T | A | C | G | | M | |
| | | 2 | 10013 | 20 | 6 | 10041 | I | 0.180 |
| | 2 | T | A | C | G | | K | 0.000 |
| | | 9919 | 2 | 16 | 9 | 9946 | T | 0.161 |
| | 3 | T | A | C | G | | R | 0.000 |
| | | 7 | 8 | 18 | 9946 | 9979 | L | 0.199 |
| | | | | | | | V | 0.000 |
| SWT5 | 1 | T | A | C | G | | M | |
| | | 1 | 10137 | 8 | 10 | 10156 | I | 0.237 |
| | 2 | T | A | C | G | | K | 0.000 |
| | | 10042 | 5 | 16 | 10 | 10073 | T | 0.159 |
| | 3 | T | A | C | G | | R | 0.000 |
| | | 10 | 5 | 24 | 10084 | 10123 | L | 0.000 |
| | | | | | | | V | 0.000 |
| SWT6 | 1 | T | A | C | G | | M | |
| | | 3 | 22556 | 38 | 17 | 22614 | I | 0.307 |
| | 2 | T | A | C | G | | K | 0.000 |
| | | 22321 | 9 | 32 | 24 | 22386 | T | 0.143 |
| | 3 | T | A | C | G | | R | 0.107 |
| | | 24 | 12 | 45 | 22378 | 22459 | L | 0.168 |
| | | | | | | | V | 0.000 |
| mix of 300 nematods | 1 | T | A | C | G | | M | |
| | | 0 | 3301 | 9 | 3 | 3313 | I | 0.629 |
| | 2 | T | A | C | G | | K | 0.152 |
| | | 3279 | 5 | 7 | 6 | 3297 | T | 0.212 |
| | 3 | T | A | C | G | | R | 0.182 |
| | | 5 | 6 | 10 | 3320 | 3341 | L | 0.272 |
| | | | | | | | V | 0.000 |

| Sample name | codon position | Phe (F) 370 | | | | | aa | % occurrence |
|---|---|---|---|---|---|---|---|---|
| WTR1 | 1 | T | A | C | G | | F | |
| | | 52042 | 35 | 47 | 58 | 52182 | L | 0.252 |
| | 2 | T | A | C | G | | V | 0.111 |
| | | 51735 | 32 | 38 | 68 | 51873 | C | 0.131 |
| | 3 | T | A | C | G | | S | 0.000 |
| | | 40529 | 63 | 10447 | 66 | 51105 | | |
| WTR2 | 1 | T | A | C | G | | F | |
| | | 38438 | 13 | 21 | 37 | 38509 | L | 0.112 |
| | 2 | T | A | C | G | | V | 0.000 |
| | | 38421 | 7 | 32 | 43 | 38503 | C | 0.112 |
| | 3 | T | A | C | G | | S | 0.000 |
| | | 20537 | 29 | 17703 | 43 | 38312 | | |
| WTR3 | 1 | T | A | C | G | | F | |
| | | 48240 | 10 | 17 | 38 | 48305 | L | 0.000 |
| | 2 | T | A | C | G | | V | 0.000 |
| | | 48485 | 9 | 21 | 52 | 48567 | C | 0.107 |
| | 3 | T | A | C | G | | S | 0.000 |
| | | 67 | 44 | 48809 | 27 | 48947 | | |
| WTR4 | 1 | T | A | C | G | | F | |
| | | 28474 | 17 | 20 | 37 | 28548 | L | 0.128 |
| | 2 | T | A | C | G | | V | 0.130 |
| | | 28374 | 15 | 24 | 33 | 28446 | C | 0.116 |
| | 3 | T | A | C | G | | S | 0.000 |
| | | 24950 | 25 | 3060 | 36 | 28071 | | |
| WTR5 | 1 | T | A | C | G | | F | |
| | | 43293 | 16 | 40 | 58 | 43407 | L | 0.000 |
| | 2 | T | A | C | G | | V | 0.134 |
| | | 43521 | 20 | 30 | 67 | 43638 | C | 0.154 |
| | 3 | T | A | C | G | | S | 0.000 |
| | | 39 | 35 | 43702 | 24 | 43800 | | |
| WTR6 | 1 | T | A | C | G | | F | |
| | | 30053 | 8 | 25 | 43 | 30129 | L | 0.170 |
| | 2 | T | A | C | G | | V | 0.143 |
| | | 29896 | 14 | 28 | 57 | 29995 | C | 0.190 |
| | 3 | T | A | C | G | | S | 0.000 |
| | | 29374 | 8 | 46 | 50 | 29478 | | |
| SWT1 | 1 | T | A | C | G | | F | |
| | | 33418 | 11 | 31 | 39 | 33499 | L | 0.134 |
| | 2 | T | A | C | G | | V | 0.116 |
| | | 33253 | 5 | 26 | 38 | 33322 | C | 0.114 |
| | 3 | T | A | C | G | | S | 0.000 |
| | | 32717 | 12 | 104 | 44 | 32877 | | |
| SWT2 | 1 | T | A | C | G | | F | |
| | | 26911 | 4 | 15 | 35 | 26965 | L | 0.120 |
| | 2 | T | A | C | G | | V | 0.130 |
| | | 27083 | 14 | 15 | 46 | 27158 | C | 0.169 |
| | 3 | T | A | C | G | | S | 0.000 |
| | | 66 | 33 | 27439 | 19 | 27557 | | |
| SWT3 | 1 | T | A | C | G | | F | |
| | | 17283 | 5 | 16 | 14 | 17318 | L | 0.000 |
| | 2 | T | A | C | G | | V | 0.000 |
| | | 17344 | 6 | 15 | 17 | 17382 | C | 0.000 |
| | 3 | T | A | C | G | | S | 0.000 |
| | | 5611 | 12 | 11724 | 15 | 17362 | | |
| SWT4 | 1 | T | A | C | G | | F | |
| | | 7548 | 3 | 5 | 16 | 7572 | L | 0.000 |
| | 2 | T | A | C | G | | V | 0.211 |
| | | 7495 | 4 | 4 | 15 | 7518 | C | 0.200 |
| | 3 | T | A | C | G | | S | 0.000 |
| | | 7385 | 2 | 26 | 7 | 7420 | | |
| SWT5 | 1 | T | A | C | G | | F | |
| | | 8379 | 8 | 8 | 12 | 8407 | L | 0.000 |
| | 2 | T | A | C | G | | V | 0.143 |
| | | 8347 | 1 | 10 | 17 | 8375 | C | 0.203 |
| | 3 | T | A | C | G | | S | 0.119 |
| | | 5869 | 5 | 2403 | 8 | 8285 | | |
| SWT6 | 1 | T | A | C | G | | F | |
| | | 18437 | 6 | 10 | 31 | 18484 | L | 0.155 |
| | 2 | T | A | C | G | | V | 0.168 |
| | | 18315 | 5 | 16 | 25 | 18361 | C | 0.136 |
| | 3 | T | A | C | G | | S | 0.000 |
| | | 17940 | 7 | 33 | 28 | 18008 | | |
| mix of 300 nematods | 1 | T | A | C | G | | F | |
| | | 3152 | 4 | 9 | 6 | 3171 | L | 0.726 |
| | 2 | T | A | C | G | | S | 0.502 |
| | | 3169 | 2 | 7 | 9 | 3187 | I | 0.126 |
| | 3 | T | A | C | G | | V | 0.189 |
| | | 1712 | 8 | 1443 | 6 | 3169 | C | 0.000 |

| Sample name | codon position | Asp (D) 371 | | | | | aa | % occurrence |
|---|---|---|---|---|---|---|---|---|
| WTR1 | 1 | T | A | C | G | | E | 0.119 |
| | | 374 | 40 | 39 | 47589 | 48042 | V | 0.322 |
| | 2 | T | A | C | G | | A | 0.203 |
| | | 155 | 47873 | 98 | 34 | 48160 | Y | 0.786 |
| | 3 | T | A | C | G | | H | 0.000 |
| | | 48502 | 44 | 48 | 58 | 48652 | D | 0.000 |
| WTR2 | 1 | T | A | C | G | | E | 0.101 |
| | | 79 | 19 | 47 | 37350 | 37495 | V | 0.000 |
| | 2 | T | A | C | G | | A | 0.350 |
| | | 33 | 37238 | 131 | 16 | 37418 | Y | 0.212 |
| | 3 | T | A | C | G | | H | 0.125 |
| | | 37622 | 6 | 40 | 38 | 37706 | D | 0.106 |
| WTR3 | 1 | T | A | C | G | | E | 0.102 |
| | | 98 | 38 | 46 | 49281 | 49463 | V | 0.000 |
| | 2 | T | A | C | G | | A | 0.330 |
| | | 34 | 48869 | 162 | 19 | 49084 | Y | 0.199 |
| | 3 | T | A | C | G | | H | 0.000 |
| | | 49003 | 17 | 52 | 50 | 49122 | D | 0.106 |
| WTR4 | 1 | T | A | C | G | | E | 0.000 |
| | | 101 | 14 | 15 | 21862 | 21992 | V | 0.295 |
| | 2 | T | A | C | G | | A | 0.231 |
| | | 65 | 21922 | 51 | 5 | 22043 | Y | 0.462 |
| | 3 | T | A | C | G | | H | 0.000 |
| | | 22162 | 16 | 33 | 22 | 22233 | D | 0.148 |
| WTR5 | 1 | T | A | C | G | | E | 0.112 |
| | | 75 | 28 | 37 | 43811 | 43951 | V | 0.000 |
| | 2 | T | A | C | G | | A | 0.287 |
| | | 29 | 43412 | 125 | 27 | 43593 | Y | 0.171 |
| | 3 | T | A | C | G | | H | 0.000 |
| | | 43483 | 10 | 33 | 49 | 43575 | D | 0.000 |
| WTR6 | 1 | T | A | C | G | | E | 0.161 |
| | | 51 | 10 | 27 | 27999 | 28087 | V | 0.110 |
| | 2 | T | A | C | G | | A | 0.333 |
| | | 31 | 28059 | 94 | 23 | 28207 | Y | 0.182 |
| | 3 | T | A | C | G | | H | 0.000 |
| | | 28513 | 6 | 36 | 46 | 28601 | D | 0.126 |
| SWT1 | 1 | T | A | C | G | | E | 0.151 |
| | | 77 | 20 | 26 | 31029 | 31152 | V | 0.125 |
| | 2 | T | A | C | G | | A | 0.371 |
| | | 39 | 31086 | 116 | 21 | 31262 | Y | 0.248 |
| | 3 | T | A | C | G | | H | 0.000 |
| | | 31594 | 10 | 32 | 48 | 31684 | D | 0.101 |
| SWT2 | 1 | T | A | C | G | | E | 0.128 |
| | | 66 | 16 | 33 | 27376 | 27491 | V | 0.102 |
| | 2 | T | A | C | G | | A | 0.303 |
| | | 28 | 27234 | 83 | 17 | 27362 | Y | 0.241 |
| | 3 | T | A | C | G | | H | 0.120 |
| | | 27299 | 15 | 22 | 35 | 27371 | D | 0.000 |
| SWT3 | 1 | T | A | C | G | | E | 0.000 |
| | | 29 | 13 | 18 | 16057 | 16117 | V | 0.000 |
| | 2 | T | A | C | G | | A | 0.337 |
| | | 14 | 15938 | 54 | 14 | 16020 | Y | 0.181 |
| | 3 | T | A | C | G | | H | 0.112 |
| | | 16013 | 5 | 17 | 15 | 16050 | D | 0.106 |
| SWT4 | 1 | T | A | C | G | | E | 0.183 |
| | | 17 | 2 | 6 | 6987 | 7012 | V | 0.000 |
| | 2 | T | A | C | G | | A | 0.412 |
| | | 5 | 6998 | 29 | 5 | 7037 | Y | 0.243 |
| | 3 | T | A | C | G | | H | 0.000 |
| | | 7091 | 0 | 9 | 13 | 7113 | D | 0.127 |
| SWT5 | 1 | T | A | C | G | | E | 0.132 |
| | | 24 | 4 | 9 | 7487 | 7524 | V | 0.159 |
| | 2 | T | A | C | G | | A | 0.332 |
| | | 12 | 7485 | 25 | 4 | 7526 | Y | 0.321 |
| | 3 | T | A | C | G | | H | 0.120 |
| | | 7568 | 2 | 6 | 10 | 7586 | D | 0.000 |
| SWT6 | 1 | T | A | C | G | | E | 0.128 |
| | | 28 | 8 | 19 | 16876 | 16931 | V | 0.000 |
| | 2 | T | A | C | G | | A | 0.294 |
| | | 14 | 16909 | 50 | 10 | 16983 | Y | 0.166 |
| | 3 | T | A | C | G | | H | 0.112 |
| | | 17141 | 5 | 18 | 22 | 17186 | D | 0.105 |
| mix of 300 nematods | 1 | T | A | C | G | | E | 0.128 |
| | | 18 | 5 | 6 | 3098 | 3127 | V | 0.256 |
| | 2 | T | A | C | G | | A | 0.576 |
| | | 8 | 3096 | 18 | 5 | 3127 | Y | 0.581 |
| | 3 | T | A | C | G | | H | 0.192 |
| | | 3111 | 2 | 3 | 4 | 3120 | D | 0.000 |
| | | | | | | | G | 0.160 |

| Sample name | codon position | Phe (F) 502 | | | | | aa | % occurrence |
|---|---|---|---|---|---|---|---|---|
| WTR1 | 1 | T | A | C | G | | F | |
| | | 42988 | 17 | 34 | 51 | 43090 | L | 0.184 |
| | 2 | T | A | C | G | | C | 0.174 |
| | | 42943 | 24 | 25 | 75 | 43067 | V | 0.118 |
| | 3 | T | A | C | G | | S | 0.000 |
| | | 437 | 79 | 42461 | 38 | 43015 | | |
| WTR2 | 1 | T | A | C | G | | F | |
| | | 28805 | 2 | 17 | 30 | 28854 | L | 0.000 |
| | 2 | T | A | C | G | | C | 0.177 |
| | | 28736 | 13 | 15 | 51 | 28815 | V | 0.104 |
| | 3 | T | A | C | G | | S | 0.000 |
| | | 22 | 21 | 28722 | 22 | 28787 | | |
| WTR3 | 1 | T | A | C | G | | F | |
| | | 39458 | 5 | 36 | 47 | 39546 | L | 0.000 |
| | 2 | T | A | C | G | | C | 0.147 |
| | | 39404 | 9 | 32 | 58 | 39503 | V | 0.119 |
| | 3 | T | A | C | G | | S | 0.000 |
| | | 25 | 33 | 39428 | 28 | 39514 | | |
| WTR4 | 1 | T | A | C | G | | F | |
| | | 24997 | 16 | 19 | 34 | 25066 | L | 0.100 |
| | 2 | T | A | C | G | | C | 0.164 |
| | | 24949 | 15 | 16 | 41 | 25021 | V | 0.136 |
| | 3 | T | A | C | G | | S | 0.000 |
| | | 94 | 25 | 24848 | 18 | 24985 | | |
| WTR5 | 1 | T | A | C | G | | F | |
| | | 36073 | 8 | 26 | 43 | 36150 | L | 0.000 |
| | 2 | T | A | C | G | | C | 0.233 |
| | | 35947 | 16 | 38 | 84 | 36085 | V | 0.119 |
| | 3 | T | A | C | G | | S | 0.105 |
| | | 22 | 28 | 36009 | 26 | 36085 | | |
| WTR6 | 1 | T | A | C | G | | F | |
| | | 24112 | 7 | 13 | 22 | 24154 | L | 0.000 |
| | 2 | T | A | C | G | | C | 0.203 |
| | | 24091 | 11 | 16 | 49 | 24167 | V | 0.000 |
| | 3 | T | A | C | G | | S | 0.000 |
| | | 18 | 14 | 24090 | 12 | 24134 | | |
| SWT1 | 1 | T | A | C | G | | F | |
| | | 31044 | 6 | 30 | 32 | 31112 | L | 0.000 |
| | 2 | T | A | C | G | | C | 0.128 |
| | | 31046 | 14 | 36 | 40 | 31136 | V | 0.103 |
| | 3 | T | A | C | G | | S | 0.116 |
| | | 17 | 21 | 31098 | 20 | 31156 | | |
| SWT2 | 1 | T | A | C | G | | F | |
| | | 24386 | 2 | 12 | 27 | 24427 | L | 0.000 |
| | 2 | T | A | C | G | | C | 0.147 |
| | | 24385 | 8 | 21 | 36 | 24450 | V | 0.111 |
| | 3 | T | A | C | G | | S | 0.000 |
| | | 20 | 6 | 24440 | 11 | 24477 | | |
| SWT3 | 1 | T | A | C | G | | F | |
| | | 15295 | 4 | 8 | 15 | 15322 | L | 0.000 |
| | 2 | T | A | C | G | | C | 0.150 |
| | | 15288 | 5 | 15 | 23 | 15331 | V | 0.000 |
| | 3 | T | A | C | G | | S | 0.000 |
| | | 11 | 14 | 15303 | 5 | 15333 | | |
| SWT4 | 1 | T | A | C | G | | F | |
| | | 6980 | 1 | 6 | 8 | 6995 | L | 0.000 |
| | 2 | T | A | C | G | | C | 0.214 |
| | | 6976 | 4 | 7 | 15 | 7002 | V | 0.114 |
| | 3 | T | A | C | G | | S | 0.000 |
| | | 3 | 5 | 7018 | 3 | 7029 | | |
| SWT5 | 1 | T | A | C | G | | F | |
| | | 7312 | 3 | 5 | 19 | 7339 | L | 0.000 |
| | 2 | T | A | C | G | | C | 0.163 |
| | | 7322 | 2 | 8 | 12 | 7344 | V | 0.259 |
| | 3 | T | A | C | G | | S | 0.000 |
| | | 6 | 5 | 7338 | 3 | 7352 | | |
| SWT6 | 1 | T | A | C | G | | F | |
| | | 14764 | 1 | 7 | 25 | 14797 | L | 0.000 |
| | 2 | T | A | C | G | | C | 0.216 |
| | | 14757 | 6 | 15 | 32 | 14810 | V | 0.169 |
| | 3 | T | A | C | G | | S | 0.101 |
| | | 10 | 7 | 14788 | 10 | 14815 | | |
| mix of 300 nematods | 1 | T | A | C | G | | F | |
| | | 2363 | 0 | 2 | 0 | 2365 | L | 0.000 |
| | 2 | T | A | C | G | | C | 0.000 |
| | | 2356 | 1 | 5 | 1 | 2363 | V | 0.000 |
| | 3 | T | A | C | G | | S | 0.212 |
| | | 2 | 0 | 2352 | 2 | 2356 | | |

| Sample name | codon position | STOP 505 | | | | | aa | % occurrence |
|---|---|---|---|---|---|---|---|---|
| WTR1 | 1 | T | A | C | G | | STOP | |
| | | 42370 | 20 | 163 | 39 | 42592 | Y | 0.383 |
| | 2 | T | A | C | G | | Q | 1.084 |
| | | 183 | 42237 | 37 | 19 | 42476 | S | 0.431 |
| | 3 | T | A | C | G | | L | 0.000 |
| | | 459 | 27 | 31 | 41813 | 42330 | E | 0.000 |
| | | | | | | | W | 0.000 |
| WTR2 | 1 | T | A | C | G | | STOP | |
| | | 28154 | 4 | 132 | 31 | 28321 | Q | 0.466 |
| | 2 | T | A | C | G | | Y | 0.153 |
| | | 13 | 28212 | 41 | 26 | 28292 | L | 0.000 |
| | 3 | T | A | C | G | | S | 0.145 |
| | | 43 | 12 | 15 | 28115 | 28185 | E | 0.109 |
| | | | | | | | W | 0.000 |
| WTR3 | 1 | T | A | C | G | | STOP | |
| | | 38315 | 9 | 186 | 36 | 38546 | Q | 0.483 |
| | 2 | T | A | C | G | | Y | 0.109 |
| | | 8 | 38431 | 48 | 18 | 38505 | L | 0.000 |
| | 3 | T | A | C | G | | S | 0.125 |
| | | 42 | 9 | 37 | 38340 | 38428 | E | 0.000 |
| | | | | | | | W | 0.000 |
| WTR4 | 1 | T | A | C | G | | STOP | |
| | | 24538 | 5 | 107 | 25 | 24675 | Q | 0.434 |
| | 2 | T | A | C | G | | Y | 0.508 |
| | | 69 | 24544 | 25 | 25 | 24663 | L | 0.280 |
| | 3 | T | A | C | G | | S | 0.101 |
| | | 125 | 7 | 15 | 24465 | 24612 | E | 0.101 |
| | | | | | | | W | 0.101 |
| WTR5 | 1 | T | A | C | G | | STOP | |
| | | 35036 | 9 | 135 | 58 | 35238 | Q | 0.383 |
| | 2 | T | A | C | G | | Y | 0.168 |
| | | 15 | 35049 | 56 | 32 | 35152 | L | 0.000 |
| | 3 | T | A | C | G | | S | 0.159 |
| | | 59 | 17 | 22 | 34997 | 35095 | E | 0.165 |
| | | | | | | | W | 0.000 |
| WTR6 | 1 | T | A | C | G | | STOP | |
| | | 23587 | 0 | 116 | 35 | 23738 | Q | 0.489 |
| | 2 | T | A | C | G | | Y | 0.144 |
| | | 5 | 23700 | 13 | 12 | 23730 | L | 0.000 |
| | 3 | T | A | C | G | | S | 0.000 |
| | | 34 | 5 | 16 | 23605 | 23644 | E | 0.147 |
| | | | | | | | W | 0.000 |
| SWT1 | 1 | T | A | C | G | | STOP | |
| | | 30631 | 6 | 134 | 26 | 30797 | Y | 0.435 |
| | 2 | T | A | C | G | | Q | 0.166 |
| | | 15 | 30653 | 30 | 20 | 30718 | S | 0.000 |
| | 3 | T | A | C | G | | L | 0.000 |
| | | 51 | 11 | 20 | 30569 | 30631 | E | 0.000 |
| | | | | | | | W | 0.000 |
| SWT2 | 1 | T | A | C | G | | STOP | |
| | | 23920 | 7 | 80 | 26 | 24033 | Y | 0.163 |
| | 2 | T | A | C | G | | Q | 0.333 |
| | | 11 | 23945 | 29 | 17 | 24002 | L | 0.000 |
| | 3 | T | A | C | G | | S | 0.121 |
| | | 39 | 2 | 17 | 23926 | 23967 | E | 0.108 |
| | | | | | | | W | 0.000 |
| SWT3 | 1 | T | A | C | G | | STOP | |
| | | 14927 | 3 | 60 | 13 | 15003 | Q | 0.400 |
| | 2 | T | A | C | G | | Y | 0.000 |
| | | 4 | 14952 | 18 | 12 | 14986 | L | 0.000 |
| | 3 | T | A | C | G | | S | 0.120 |
| | | 10 | 6 | 11 | 14908 | 14924 | E | 0.000 |
| | | | | | | | W | 0.000 |
| SWT4 | 1 | T | A | C | G | | STOP | |
| | | 6944 | 2 | 28 | 6 | 6980 | Q | 0.401 |
| | 2 | T | A | C | G | | Y | 0.130 |
| | | 3 | 6927 | 8 | 6 | 6944 | L | 0.000 |
| | 3 | T | A | C | G | | S | 0.115 |
| | | 9 | 2 | 5 | 6920 | 6931 | E | 0.000 |
| | | | | | | | W | 0.000 |
| SWT5 | 1 | T | A | C | G | | STOP | |
| | | 7220 | 2 | 24 | 6 | 7252 | Q | 0.331 |
| | 2 | T | A | C | G | | Y | 0.249 |
| | | 1 | 7226 | 9 | 3 | 7239 | L | 0.000 |
| | 3 | T | A | C | G | | S | 0.124 |
| | | 10 | 1 | 8 | 7219 | 7230 | E | 0.000 |
| | | | | | | | W | 0.000 |
| SWT6 | 1 | T | A | C | G | | STOP | |
| | | 14565 | 3 | 60 | 18 | 14646 | Q | 0.410 |
| | 2 | T | A | C | G | | Y | 0.117 |
| | | 1 | 14582 | 13 | 18 | 14614 | L | 0.000 |
| | 3 | T | A | C | G | | S | 0.000 |
| | | 17 | 8 | 10 | 14527 | 14552 | E | 0.123 |
| | | | | | | | W | 0.123 |
| mix of 300 nematods | 1 | T | A | C | G | | STOP | |
| | | 2272 | 1 | 5 | 0 | 2278 | Y | 0.142 |
| | 2 | T | A | C | G | | Q | 0.219 |
| | | 1 | 2208 | 0 | 1 | 2210 | L | 0.000 |
| | 3 | T | A | C | G | | S | 0.000 |
| | | 3 | 0 | 0 | 2115 | 2118 | E | 0.000 |
| | | | | | | | W | 0.000 |

major nucleic acid
